# Eosinophils restrict CRC metastasis by inhibiting pro-tumorigenic SPP1^+^ macrophage differentiation

**DOI:** 10.1101/2024.12.13.628333

**Authors:** Kristina Handler, Deeksha Raju, Alessandra Gurtner, Tosca Dalessi, Michael D. Brügger, Daniel Crepaz, Cinzia Esposito, Zlatko Trajanoski, Christian Münz, Michael Scharl, Tomas Valenta, Salvatore Piscuoglio, Hassan Fazilaty, Isabelle C. Arnold

**Author notes:** These authors contributed equally to this work (Kristina Handler, Deeksha Raju, Alessandra Gurtner). Dimericon Therapeutics AG, Bachtobelstrasse 5, 8810 Horgen.

## Abstract

Eosinophils, traditionally associated with allergic responses, have emerged as critical immune modulators in colorectal cancer (CRC). Here, we reveal that eosinophils actively shape the tumor microenvironment and influence metastatic progression. Using comprehensive transcriptomics analysis of human CRC and a murine orthotopic tumor model, we identify a conserved tumor-specific eosinophil signature and activation profile. Despite their declining presence in advanced CRC, eosinophils suppress metastatic dissemination by counteracting the pro-tumorigenic functions of SPP1^+^ macrophages – a subset linked to immune exclusion and tumor metastasis. Mechanistically, eosinophils respond to tumor-derived signals and inhibit macrophage differentiation into SPP1^+^ cells. Eosinophil depletion exacerbates peritoneal tumor spread. These findings highlight the pivotal role of eosinophils in restraining late-stage CRC progression and unveil a novel eosinophil–macrophage axis as potential therapeutic targets.

## Introduction

Colorectal cancer (CRC) is a leading cause of cancer-related mortality worldwide, primarily due to tumor metastasis to distant sites such as the liver, lungs, and peritoneal cavity (*1*). Among these, peritoneal metastases are associated with the poorest prognosis (*2*, *3*). Despite advances in early detection and treatment, late-stage CRC remains largely refractory to immunotherapies, especially in subtypes characterized by a low mutational burden, lack of neoantigens, and poor immunoreactivity (*4*). This underscores the urgent need to better understand and target the tumor immune landscape.

The tumor microenvironment (TME) consists of a variety of cells that can modulate tumor growth, immune evasion, and metastasis. Among these, eosinophils – a granulocyte subset enriched in mucosal tissues – have recently emerged as intriguing orchestrators of immune responses and tissue homeostasis (*5, 6*). While traditionally associated with allergic diseases, eosinophils are increasingly recognized for their roles in tissue remodeling, immune regulation and anti-tumor immunity. Notably, eosinophil infiltration is often associated with improved prognosis in gastrointestinal malignancies (*7*), including CRC, and clinical evidence links their presence to enhanced responses to immune checkpoint blockade therapies (*8*–*10*). However, their precise functions and interactions within the local immune network, particularly during CRC progression and metastasis, remain poorly understood.

Recent studies have identified SPP1^+^ tumor-associated macrophages (TAMs) as key drivers of metastasis and immune exclusion in CRC (*11*). These macrophages are known to interact with cancer-associated fibroblasts to create a tumor-supportive niche (*12*). However, whether they also interact with eosinophils and how such interactions might impact metastasis and the tumor immune network remains unexplored.

Here, we present a comprehensive investigation of eosinophil dynamics within the TME in human CRC and a murine CRC model. Our findings reveal conserved eosinophil transcriptional programs across species, highlighting their activation and functional reprogramming in tumors. Despite a marked decline in eosinophil frequencies with advancing CRC stages, their activation state increases in more malignant stages, suggesting stage-specific roles in tumor progression. Notably, eosinophils suppress peritoneal metastases by modulating the differentiation of pro-tumorigenic SPP1^+^ TAMs. Mechanistically, we show that eosinophils respond to tumor-derived signals, adopt an activated phenotype and prevent SPP1^+^ TAMs differentiation. Together, these findings establish a previously unrecognized eosinophil–macrophage axis in CRC progression and metastasis, offering new avenues for therapeutic interventions.

## Results

### Eosinophils are highly activated across progressive CRC stages in patients

To characterize eosinophil dynamics in progressive CRC stages, we quantified MBP-1^+^ eosinophils using immunofluorescence staining on human tissue microarrays. In line with our previous findings (*13*), eosinophil frequencies decreased markedly as CRC progressed from stage I to stage III (**Fig. 1A**). Notably, their expression of PD-L1 – a marker that we have previously associated with an activated eosinophil subset known as A-Eos (*14*) – showed a dichotomic pattern, with lower A-Eos frequencies found in precancerous adenoma stages and higher frequencies observed across all adenocarcinoma stages compared to normal colonic tissue (**Fig 1B, C and fig. S1A**).

**Fig 1.**
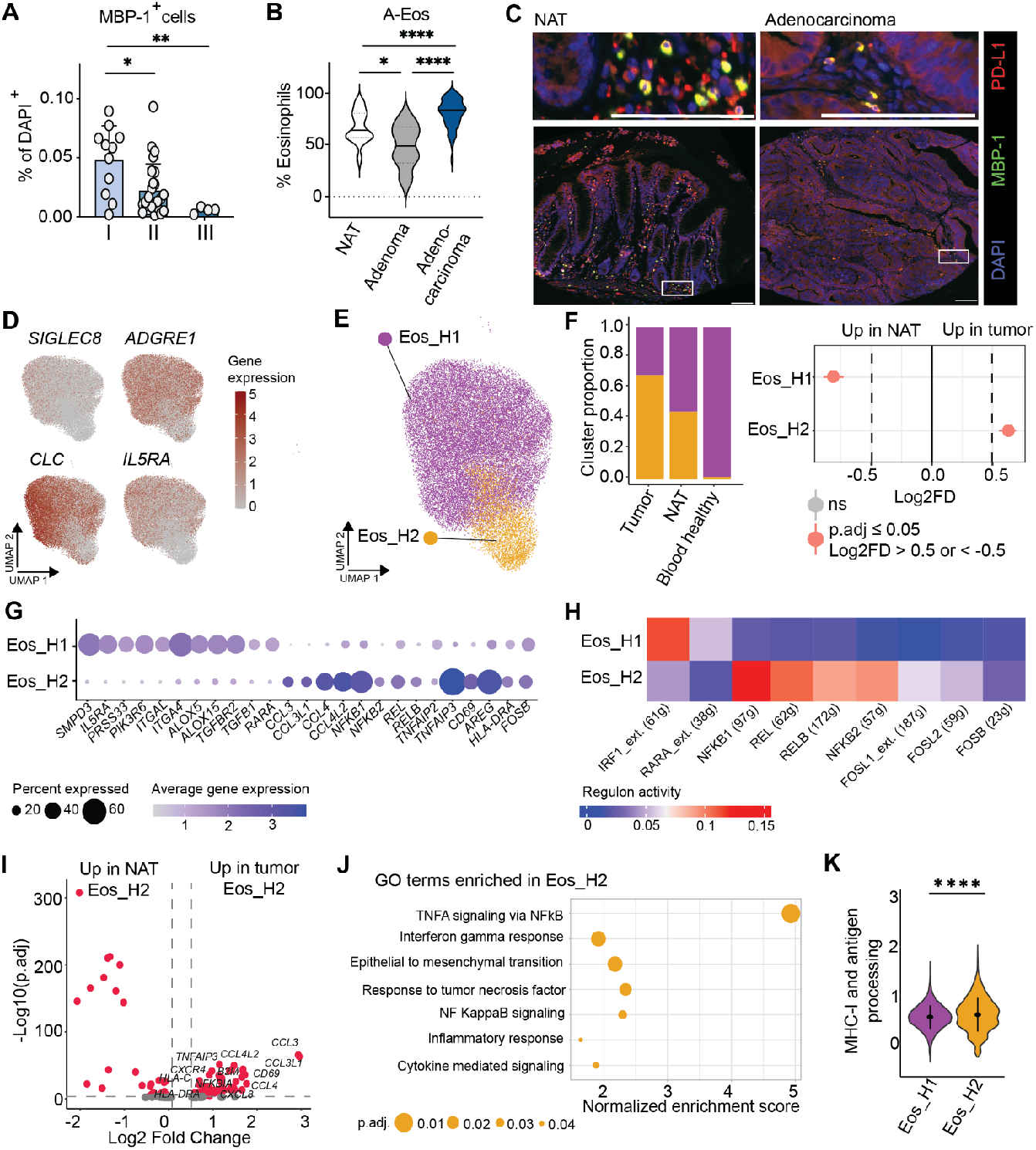
Characterisation of human eosinophils in colorectal cancer at single cell resolution. (**A**) MBP-1^+^ eosinophils as a proportion of DAPI^+^ cells on human tissue microarrays, across increasing CRC stages (I n = 11, II n = 25, III n = 4 slides). (**B**) A-Eos frequencies (co-localization of MBP-1^+^ and PD-L1^+^ signal) in normal adjacent tissue (NAT), adenoma and adenocarcinoma on tissue microarrays (NAT n = 24, adenoma n = 6, adenocarcinoma n = 28 slides). (**C**) Immunofluorescence images of tissue microarrays from NAT (left) and adenocarcinoma (right). Eosinophils are represented in green (stained for MBP-1), with PD-L1 expression in red and DAPI in blue. Scale bar: 100 µm. (**D** and **E**) UMAP of the integrated tumor, NAT and blood healthy eosinophils. The color represents the gene expression level of selected marker genes (D) and different clusters (E). (**F**) Barplot (left) indicating proportions of clusters within different tissues. Statistical analysis between tumor and NAT shown in a point-range plot (right) [Permutation test, Benjamini Hochberg adjusted p-values, ns (non significant), Log2FD (log2 fold difference)]. (**G**) Bubble plot displaying the average gene expression of selected DEGs (differentially expressed genes) per cluster. All genes are significant (p.adj ≤ 0.05, Log2 Fold Change > 0.25 or < -0.25). (**H**) Heatmap showing the regulon activity of selected regulons from SCENIC analysis of each cluster. (**I**) Volcano plot of DEGs between NAT and tumor eosinophils from cluster Eos_H2. Red dots represent significantly up- and down-regulated genes (p.adj ≤ 0.05, Log2 Fold Change > 0.25 or < -0.25). Selected genes are highlighted. (**J**) Bubble plot indicating enriched GO (gene ontology) terms in Eos_H2 compared to the Eos_H1. The x-axis displays the normalized enrichment score and the size shows p.adj (null hypothesis testing and Benjamini-Hochberg correction). (**K**) Violin plot depicting the MHC-I and antigen processing score between Eos_H1 and Eos_H2 eosinophils (two-sided Wilcoxon Rank-Sum test). (A, B) One-Way ANOVA. (A, B, K) Mean ± standard deviation are indicated; stars indicate significance of the p-value (* < 0.05, ** < 0.01, *** < 0.001, **** < 0.0001). (D-K) Tumor and NAT n = 7, blood healthy n = 6 samples. (G, I) DEG analysis: non-parametric Wilcoxon Rank-Sum Test and Bonferroni correction of p-values. (B-K) NAT: normal adjacent tissue. (F, G, I, J) P.adj: p-value adjusted.

To examine the transcriptional subset composition of human eosinophils in CRC, we performed single-cell RNA sequencing (scRNA-seq) on CD45^+^ leukocytes, enriched from tumors and matched normal adjacent tissue (NAT) biopsies from seven CRC patients (**Fig. S1B and table S1**). Blood eosinophils from six healthy individuals were included as controls. Eosinophil clusters were identified based on the expression of specific marker genes, including *CLC, IL5RA, ADGRE1* and *SIGLEC8* (**Fig. 1D**) and validated using a mouse eosinophil-specific gene signature, which showed the strongest signal in the annotated eosinophil clusters (**fig. S1C**). Eosinophils from all three sources were integrated (**fig. S1D**). Their quality was consistently high across tissues and clusters (**fig. S1E, F**). Clustering analysis identified two main eosinophil subsets, Eos_H1 and Eos_H2, present in variable proportions across tissues and patients (**Fig. 1E, F and fig. S1H**). Notably tumor eosinophils predominantly shifted towards the Eos_H2 cluster compared to eosinophils from normal adjacent tissue, suggesting enrichment of this cluster in CRC (**Fig. 1F)**.

Differential gene expression and regulon activity (SCENIC) analysis revealed distinct characteristics for each subset (**Fig. 1G, H**). The Eos_H1 cluster expressed multiple genes previously reported in human peripheral blood eosinophils, such as *SMPD3, PRSS33, PIK3R6* and *IL5RA (15)*, consistent with its predominant representation among blood eosinophils (**Fig. 1F**). This subset also showed enrichment for cell adhesion-related genes (*ITGAL, ADGRG5*) and for genes regulated by the transcription factor Interferon Regulatory Factor 1 (IRF1). In contrast, Eos_H2 upregulated the eosinophil activation marker *CD69 (16)* and genes associated with immune regulation and inflammation, including *CCL3, CCL4*, and *TNFAIP3*. Eos_H2 also exhibited strong activation of transcription factors from the NF-κB pathway (*17*) (*REL, RELB, NFKB1, NFKB2*) and AP-1 complex (*FOSL1, FOSL2, FOSB*) (*18*) (**Fig. 1G, H**). Together, these results highlight the dynamic changes in eosinophil phenotypes during CRC progression and emphasize their activation via the NF-κB pathway within the TME.

### Human eosinophils display tumor-specific gene signatures distinct from those of adjacent colonic tissue

To assess whether eosinophils exhibit distinct gene expression profiles within the tumor core compared to normal adjacent tissue, we conducted differential gene expression analysis combined with gene ontology (GO) analysis. Intratumoral eosinophils displayed marked transcriptional differences compared to their counterparts in non-tumor tissue (**Table. S2**), with most upregulated genes associated with the Eos_H2 cluster (**fig. S1G**). Notably, these upregulated genes primarily included cytokines and chemokines (e.g., *CCL3, CCL3L1, CCL4, CCL4L1, CXCL8, CXCR4*), NF-κB signaling (*NFKBIA*), TNF-alpha signaling (*TNFAIP3*), and MHC-I components (*HLA-C, HLA-DRA, B2M*) (**Fig. 1I**). GO analysis (**Fig. 1J**) and elevated scores for MHC-I and antigen processing signatures (**Fig. 1K**) further showed enrichment in these pathways, indicating enhanced immunoregulatory activity. These findings suggest that eosinophils undergo functional reprogramming in the TME, with the Eos_H2 subset representing an activated, inflammation-associated phenotype.

### Mouse eosinophils recapitulate human eosinophil recruitment patterns and transcriptional profiles in an orthotopic model of late-stage CRC

To explore the functional role of eosinophils in advanced CRC stages, we employed an orthotopic mouse model of tumor organoid transplantation. AKPS colonic tumor organoids – harboring the four most common CRC-driving mutations [*Apc*-LOF (loss of function), *p53*-LOF, *Kras*-GOF (gain of function), *Smad4*-LOF] (*19*) – were successfully engrafted into the colonic mucosa of C57BL/6JRj mice following colonoscopy-guided injection. Within six weeks, tumors developed and progressively expanded through the colonic layers toward the peritoneal cavity (**Fig. 2A**), mirroring some of the invasive characteristics of late-stage human CRC. Intriguingly, eosinophil infiltration was significantly elevated in precancerous lesions and at an earlier time point (4 weeks) of AKPS tumor development compared to normal adjacent tissue, but stabilized at later stages (6 weeks), where eosinophil numbers were comparable between tumors and normal adjacent tissue (**Fig. 2B, fig. S2A**). Despite this plateau, eosinophil activation markedly increased during malignant transformation, with a higher proportion of activated eosinophils (A-Eos) in advanced AKPS tumors (**Fig 2C and fig. S2B**). This contrasts with findings in *Apc*^Min/+^ adenomas (*20*), where eosinophils were more abundant but predominantly consisted of the less activated CD80^−^ PD-L1^−^ B-Eos subtype (*14*) (**Fig. 2C, D and fig. S2B)**. Overall, these findings align with our observations in CRC patients, where eosinophil frequencies increased in adenomas but decreased in malignant stages while displaying an enhanced activation profile (**Fig. 1A-C**).

**Fig 2.**
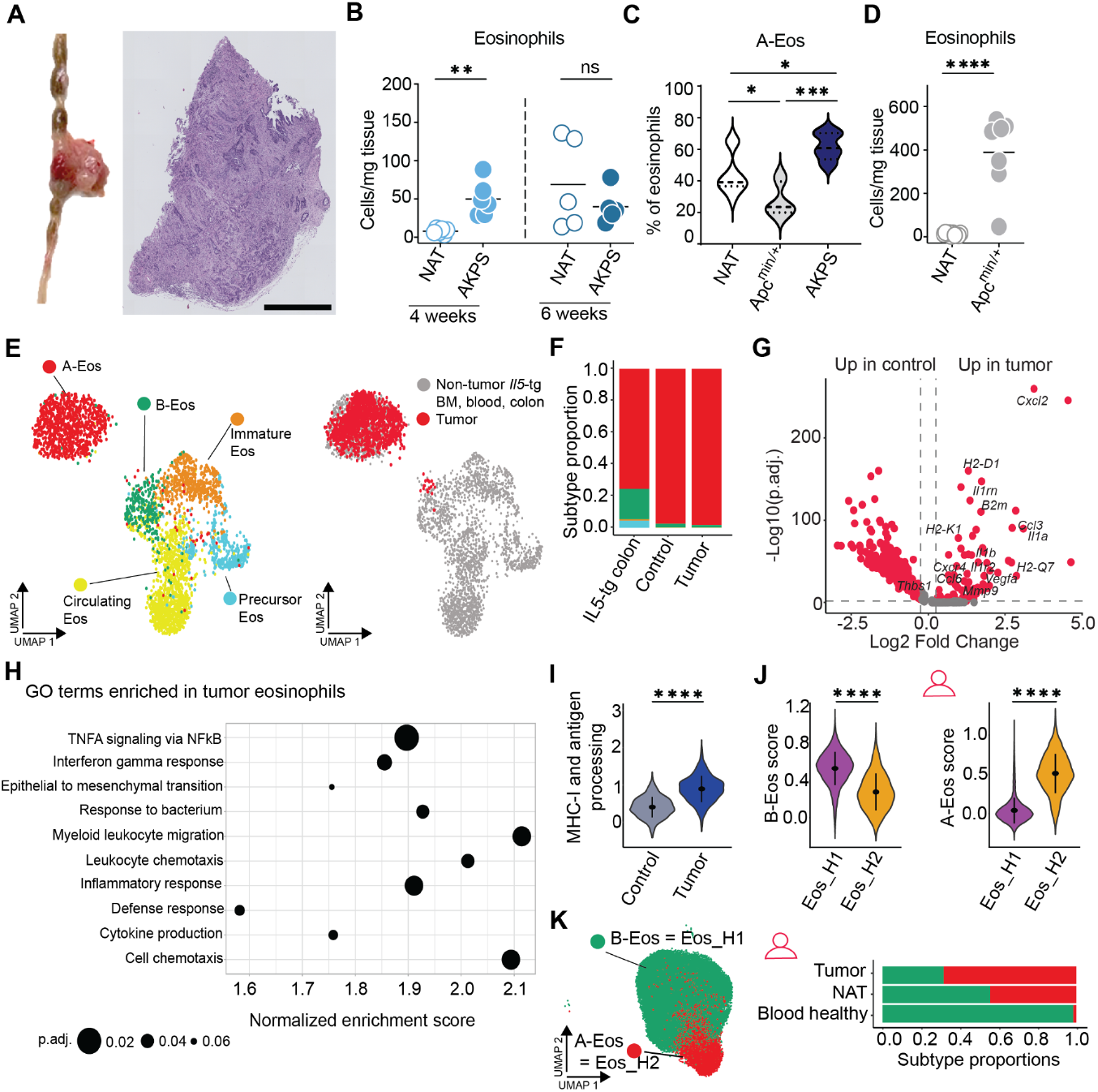
Mouse colorectal cancer eosinophils and their comparison to the human counterparts. (**A**) Macroscopic (left) and histological H&E (hematoxylin and eosin) (right) image of the AKPS tumor. Scale bar: 1 mm. (**B**) Absolute counts of eosinophils in normal adjacent tissue (NAT) and AKPS tumors after 4 and 6 weeks (NAT and AKPS 4 weeks n = 6, NAT and AKPS 6 weeks n = 5 samples). (**C**) A-Eos frequencies in NAT, Apc^min/+^, and AKPS tumors (NAT n = 13, Apc^min/+^ n = 7, AKPS tumor n = 5 samples). (**D**) Absolute counts of eosinophils in NAT and Apc^min/+^ (NAT n = 19, Apc^min/+^ n = 7 samples). Unpaired t-test. (**E**) UMAP of integrated tumor eosinophils with eosinophils from non-tumor bearing *Il5-*tg (*Il5-*transgenic) BM (bone marrow), blood and colon (*14*). Colors represent eosinophil subtypes (left) and tumor derived eosinophils (right). (**F**) Eosinophils subtype proportions between different conditions. (**G**) Volcano plot depicting DEGs (differentially expressed genes) between eosinophils from control and tumor. Significantly enriched or depleted genes are highlighted in red (p-value adjusted ≤ 0.05, Log2 Fold Change > 0.25 or < -0.25, non-parametric Wilcoxon Rank-Sum test, Bonferroni correction of p-values). Selected DEGs are highlighted. (**H**) Bubble plot indicating enriched GO (gene ontology) terms in tumor eosinophils compared to the control. The x-axis displays the normalized enrichment score and the size shows the adjusted p-value (p.adj., null hypothesis testing and Benjamini-Hochberg correction. (**I**) Violin plot of the MHC-I and antigen processing score between control and tumor eosinophils. (**J**) Violin plots highlighting mouse eosinophilic subtype scores (left: B-Eos, right: A-Eos) within human eosinophil clusters. (**K**) UMAP of human eosinophils colored by annotated subtypes (left) and a barplot showing their proportions within different tissues (right). (B,C) One-Way ANOVA. (B-D, I, J) Mean ± standard deviation are indicated; stars indicate significance of the p-value (* < 0.05, ** < 0.01, *** < 0.001, **** < 0.0001, ns non-significant). (E-I) Control and tumor n = 5 samples. (I, J) Two-sided Wilcoxon Rank-Sum test. (J, K) Tumor and NAT n = 7, blood healthy n = 6 samples. (B-D, J, K) NAT: normal adjacent tissue.

To compare the transcriptional profiles of mouse and human CRC eosinophils, we performed scRNA-seq on CD45-enriched leukocytes isolated from AKPS tumors six weeks post-injection. Colons and blood of naïve (control) mice were further included for comparison.

The quality of extracted eosinophils was consistent across samples (**fig. S2C**). Newly sequenced cells were projected onto our recently published reference of eosinophil subtypes (*14*) using a label transfer approach (**fig. S2D**). Most tumor eosinophils mapped to the A-Eos cluster (**Fig. 2E)**. Of note, eosinophils isolated from the colon and blood of wild-type mice displayed a more differentiated profile compared to those previously reported in *Il5*-tg mice (*14*) (**Fig. 2F and fig. S2E)**, a strain with elevated eosinophil numbers across tissues (*21*). Similar to our observations in human CRC, intratumoral eosinophils upregulated multiple transcripts associated with cytokine and chemokine signaling (e.g., *Ccl3, Cxcr4, Il1a, Il1rn, Il1r2, Ccl6*), MHC-I components (*H2-Q7, H2-K1, B2m, H2-D1*), and tissue remodeling (*Vegfa, Mmp9, Thbs1*) (**Fig. 2G**). These transcriptional changes were further reflected by the enrichment of GO terms such as “Epithelial to mesenchymal transition, Inflammatory response, Interferon gamma response and TNFα signaling via NF-κB”, along with an enhanced “MHC-I and antigen processing” signature score compared to control colon (**Fig. 2H, I**).

To evaluate transcriptional similarities between mouse and human eosinophils and assess potential correspondence between their respective subpopulations, we determined the top differentially expressed genes (DEGs) for each mouse subset and calculated signature scores for human eosinophil clusters. This analysis mapped human Eos_H1 to the mouse B-Eos subset and human Eos_H2 to the mouse A-Eos subset (**Fig. 2J)**, confirming the enrichment of A-Eos in CRC patient tumors (**Fig. 2K**). Comparison of eosinophil transcripts further revealed interspecies similarities ranging from 23% to 45% across tissues, with the strongest correspondence observed in the A-Eos subset (**fig. S2F, G**). Together, these findings underscore the high concordance between human and mouse eosinophil subsets and highlight the conserved gene expression programs of activated eosinophils within the TME.

### Neutrophils and SPP1^+^ TAMs accumulate in the TME of human and mouse CRC

To further examine the TME of human and mouse CRC, we compared the immune cell composition of tumors and normal adjacent or naive tissue in the respective scRNA-seq dataset from each species. This analysis revealed notable differences in cellular proportions between tumors and control tissues (**Fig. 3A, B and fig. S3A-C**). Specifically, eosinophils, mature B cells and plasma cells were reduced within tumors, while neutrophils and SPP1^+^ TAMs were significantly increased in both species (**Fig. 3C and fig. S3B**). SPP1^+^ TAMs were distinguished from other monocytes and macrophages by their high expression of the *SPP1/Spp1* gene (**fig. S3A**). Along with C1QC^+^ TAMs, these cells are thought to originate from monocyte-like FCN1^+^ cells (*22*, *23*). In the human dataset, monocytes corresponded to the FCN1^+^ monocyte-like population (characterized by high *FCN1* expression), while TAMs aligned with the C1QC^+^ (high *C1QC, C1QB*) and SPP1^+^ (high *SPP1*) subsets. Similarly, in the mouse dataset, we identified C1QC^+^ and SPP1^+^ TAM populations, although the monocytes showed transcriptional differences, such as low expression of *Fcna* (**fig. S3A**).

**Fig 3.**
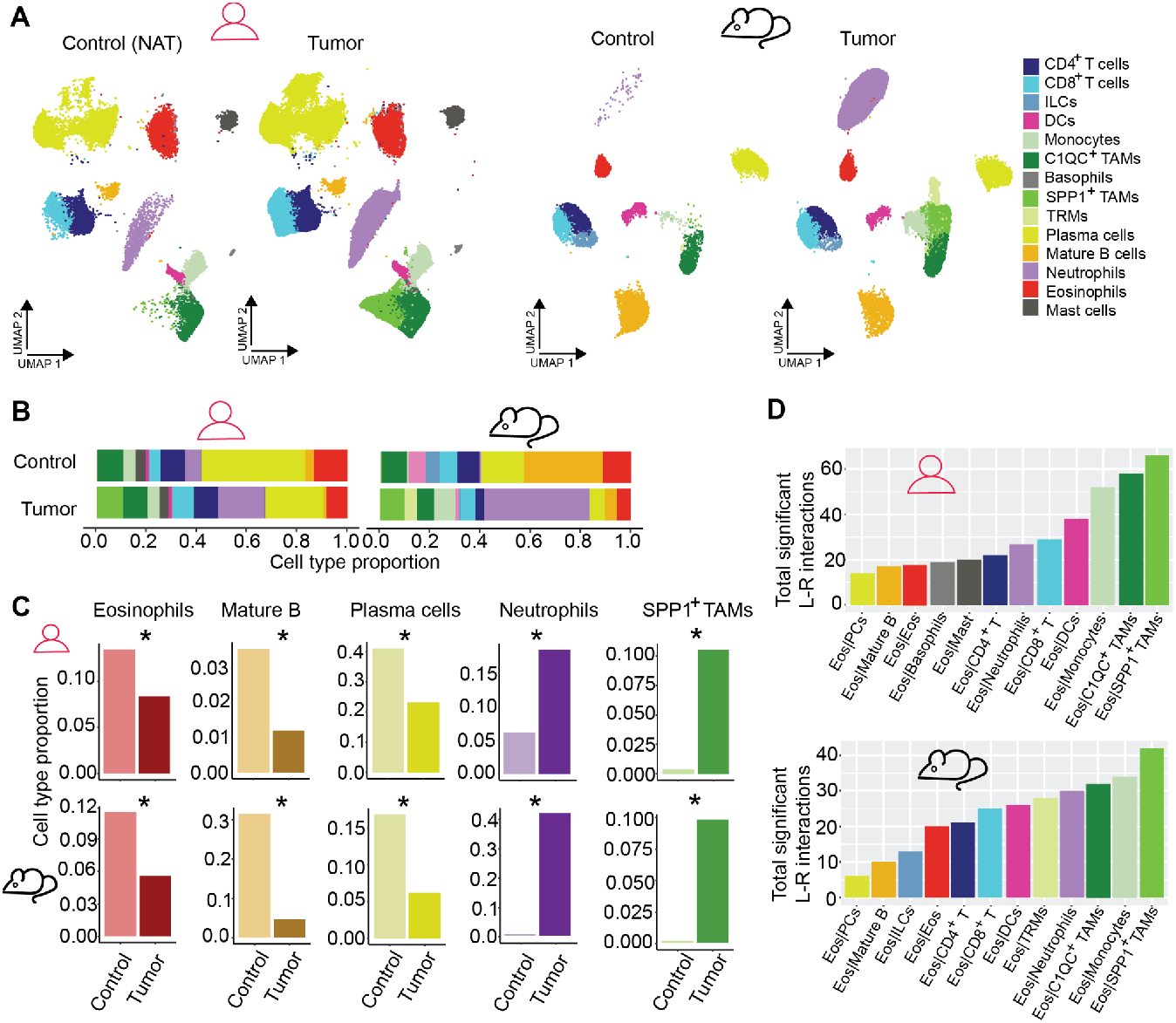
Analysis of immune cells from the tumor immune network in human and mouse colorectal cancer. (**A**) UMAP of CD45^+^ cell type populations from control and tumor of human (left) and mouse (right) data. Colors represent annotated cell types. (**B**) Barplots comparing cell type proportions from human (left) and mouse (right) data. NAT: normal adjacent tissue. (**C**) Barplots of selected cell types comparing control and tumor of human (left) and mouse (right) datasets. Stars indicate statistical significance (adjusted p-value ≤ 0.05 and Log2 Fold Difference > 0.5 or < -0.5, permutation test, Benjamini-Hochberg p-value adjustment). (**D**) Total number of significant L-R (ligand-receptor) interactions between eosinophils and any other cell type within tumors (upper plot: human; lower plot: mouse). L-R interactions were predicted using the CellPhoneDB algorithm and database (null hypothesis testing, p-value ≤ 0.05). Both directionalities (ligand-receptor and receptor-ligand) were considered. (A-D) TAMs: Tumor associated macrophages, B: B cells, T: T cells, Eos: Eosinophils, PCs: Plasma cells, DCs: Dendritic cells, ILCs: Innate lymphoid cells, TRMs: Tissue resident macrophages. Tumor and NAT from human datasets n = 7, tumor and control from mouse datasets n = 5 samples.

To explore eosinophil interactions with immune cells in the TME, we performed ligand-receptor prediction analysis using CellPhoneDB (*24*). Significant ligand-receptor interactions were quantified, accounting for both eosinophils acting as ligands and as receptors. Interestingly, in both species, eosinophils exhibited the highest number of predicted interactions with the monocyte/macrophage compartment, particularly with SPP1^+^ TAMs (**Fig. 3D and fig. S3D**). These observations suggest that eosinophil-macrophage interactions may have functional relevance within the TME.

### Eosinophil deficiency promotes tumor metastasis and drives the expansion of SPP1^+^ TAMs

To examine the impact of eosinophils on late-stage tumor development and define underlying mechanisms, we injected AKPS organoids into the colon of PHIL mice, a strain genetically engineered to lack mature eosinophils through the expression of diphtheria toxin A under the eosinophil peroxidase (*Epx*) promoter (*25*). Six weeks after organoid transplantation, PHIL mice developed smaller primary tumors compared to their wild-type (WT) littermates, with significantly reduced tumor weight and volume (**Fig. 4A, B**). However, PHIL mice exhibited a marked increase in both the incidence and number of peritoneal tumor dissemination compared to WT mice (**Fig. 4C, D**), a manifestation of advanced disease association with particularly poor prognosis in patients (*2*, *3*). The scRNA-seq profiling of CD45^+^ immune cells from primary and peritoneal tumors in PHIL and WT mice revealed a progressive increase in SPP1^+^ TAMs – and a corresponding decrease in eosinophils – from WT control tissues to tumors and ultimately to peritoneal metastases (**Fig. 4E and fig. S4B**). In PHIL mice, eosinophil deficiency resulted in a further expansion of SPP1^+^ TAMs within tumors, while other macrophage and monocyte subsets were either unaffected or decreased (**Fig. 4E and fig. S4C**).

**Fig 4.**
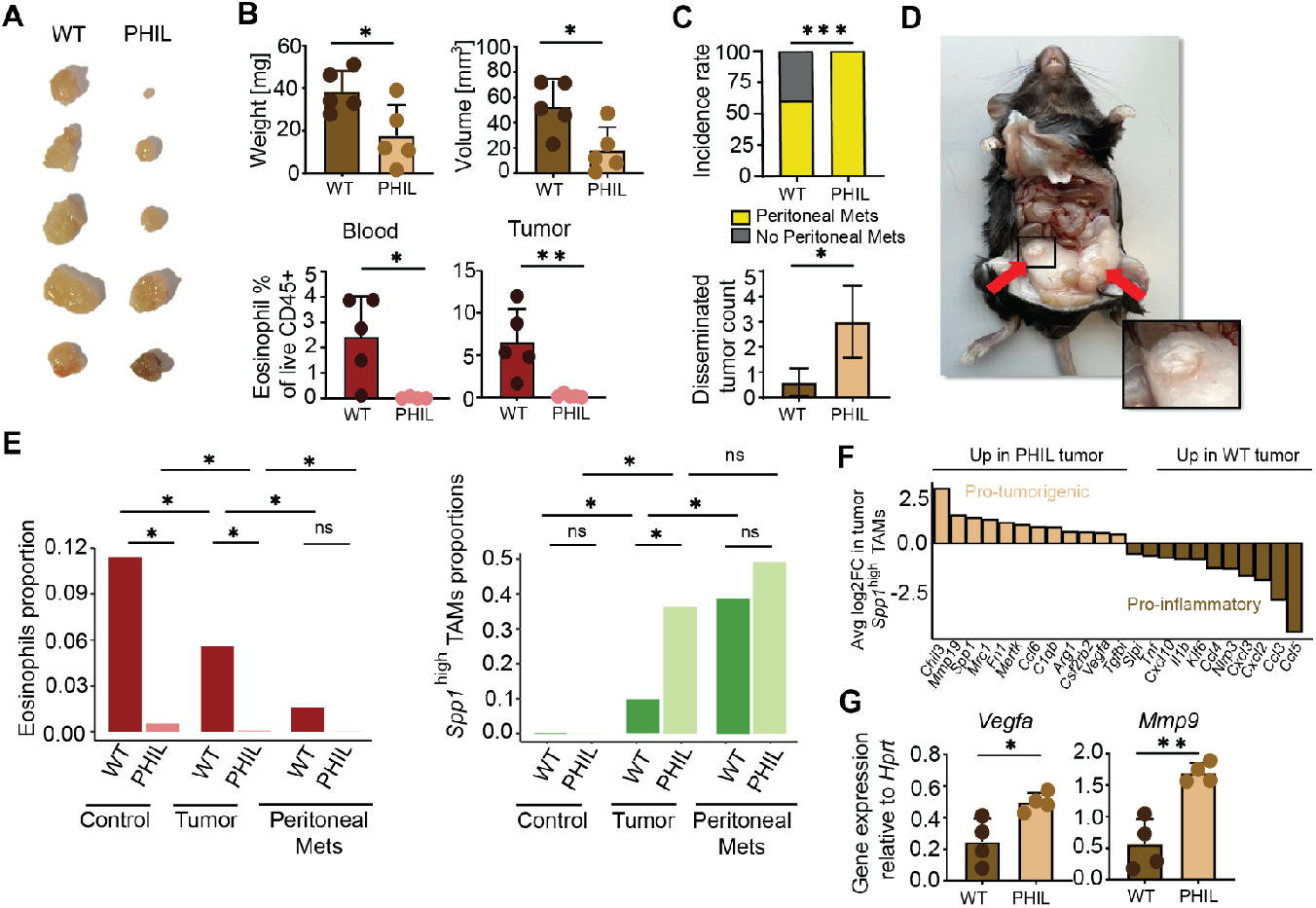
Eosinophil deficiency drives tumor metastasis and the expansion of SPP1^+^ TAMs. (**A**) Macroscopic images of AKPS primary tumors from WT and PHIL mice, taken at the endpoint of 6 weeks. (**B**) Tumor weights (upper left) and volumes (upper right), percentage of eosinophils in the blood (bottom left) and tumor (bottom right) from AKPS tumor-bearing mice (n = 5 samples each). (**C**) The incidence rate (upper) and counts (bottom) of peritoneal metastasis in mice bearing primary AKPS tumors (WT n = 4, PHIL n = 2). Mets: metastasis. Two-tailed Fisher’s exact tests (upper plot). (**D**) Image of a PHIL mouse injected with AKPS organoids. Red arrows point to metastasis in the peritoneal cavity. (**E**) Bar plots of eosinophils (left) and SPP1^+^ TAMs (right) comparing the proportions between WT and PHIL from different tissues. Stars indicate statistical significance [adjusted p-value ≤ 0.05 and Log2 Fold Difference > 0.5, ns (non-significant), permutation test, Benjamini-Hochberg p-value adjustment]. WT control and tumor n = 5, PHIL control and tumor n = 2, WT peritoneal mets = 3, PHIL peritoneal mets = 6 samples. (**F**) Selected DEGs (differentially expressed genes) between SPP1^+^ TAMs from PHIL and WT tumors. The Average log2 fold change (Avg. log2FC) is indicated on the y-axis. Genes are all significant (p-value adjusted ≤ 0.05); non-parametric Wilcoxon Rank-Sum test and Bonferroni correction of p-values. WT control and tumor n = 5, PHIL control and tumor n = 2 samples. (**G**) Relative gene expression to house-keeping gene *Hprt* of *Vegfa* and *Mmp9* in whole AKPS tumor tissue from WT vs PHIL mice, as determined by RT-qPCR (n = 4 samples each). (B, C bottom) Unpaired t-test, mean ± standard deviation is indicated. (B, C, E, G) stars indicate significance of the p-value (* < 0.05, ** < 0.01, *** < 0.001, **** < 0.0001).

Beyond these quantitative changes, SPP1^+^ TAMs in PHIL tumors upregulated multiple transcripts linked to pro-tumorigenic activities (**Fig. 4F**). Consistently, bulk expression of genes associated with angiogenesis (*Vegfa*) and tissue remodeling (*Mmp9*) was significantly enriched in tumors from PHIL mice (**Fig. 4G**).

Together, these findings suggest that, while eosinophils do not limit primary tumor growth in late-stage CRC, they play a critical role in preventing metastatic tumor dissemination by restraining the expansion and pro-tumorigenic activities of SPP1^+^ TAMs.

### Eosinophils are activated by AKPS tumor cells and restrict the differentiation of SPP1^+^ macrophages in vitro

Our findings indicate that both murine and human eosinophils are highly activated in the TME of advanced CRC stages (**Fig. 1B-F and fig. 2C-E**). To determine if this activation is influenced by tumor cell-derived factors, we co-cultured eosinophils purified from the spleens of *Il5*-tg mice with either dissociated AKP or AKPS organoids. AKP organoids lack the *Smad4* loss-of-function mutation present in AKPS and represent a less advanced CRC stage. Notably, co-culture with both organoid types shifted eosinophil maturation toward the (CD80^+^ PD-L1^+^) A-Eos subset, with higher proportions and increased Siglec-F expression in the AKPS condition (**Fig. 5A and fig. S5A**). This activation was accompanied by increased surface expression of CD11b, β2 integrin and CD63, along with decreased granularity (SSC-A), suggesting enhanced adhesion and degranulation (**Fig. 5B, C**). Remarkably, these features were recapitulated when eosinophils were cultured in the presence of AKPS-derived supernatant (AKPS-sup), indicating that eosinophil activation is driven by soluble tumor-derived factors (**fig. S5B, C**).

**Fig 5.**
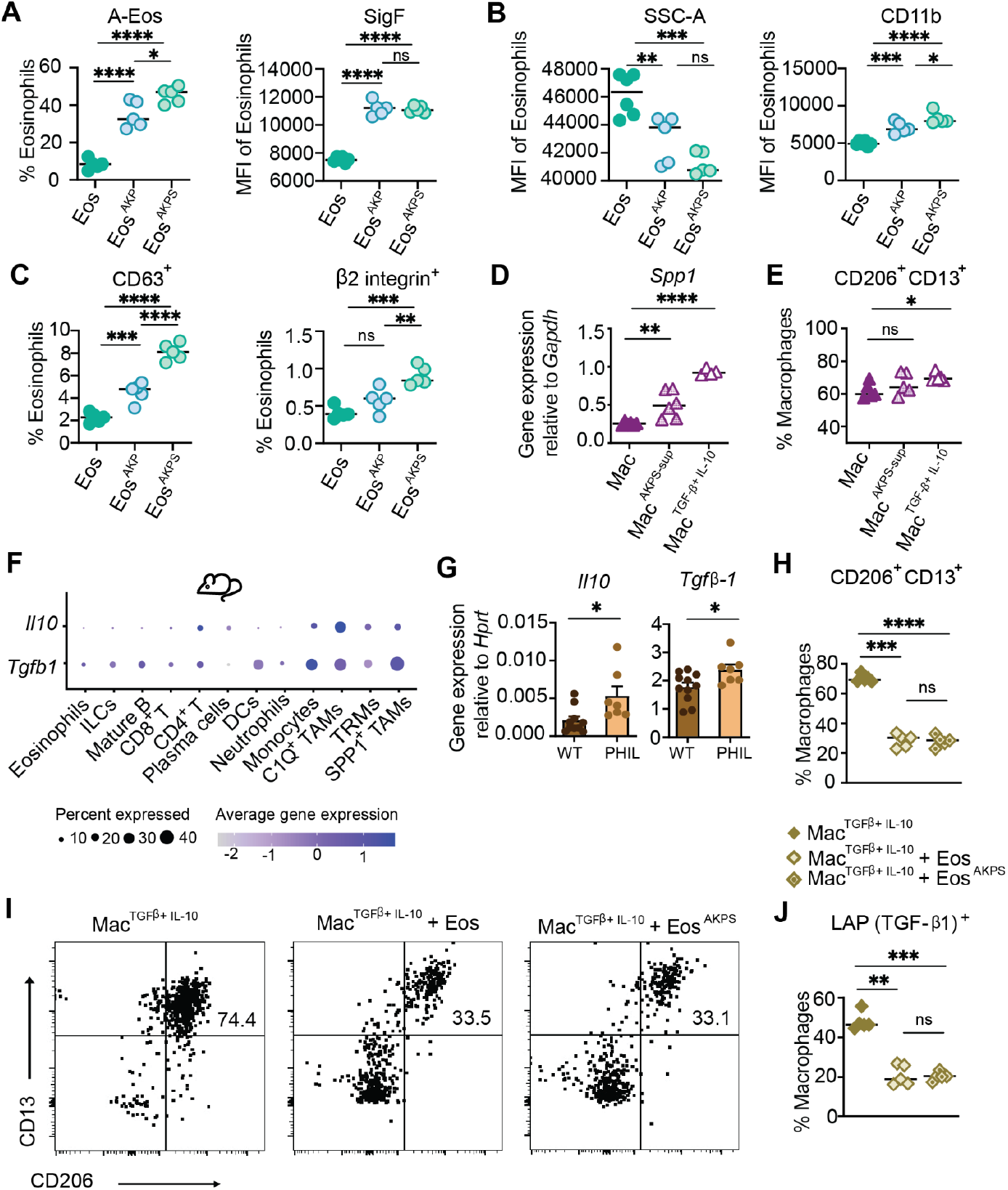
Eosinophils restrict the differentiation of SPP1^+^ macrophages *in vitro*. (**A**-**C**) A-Eos proportion (A left), Mean Fluorescence Intensity (MFI) of Siglec-F (SigF) (A right), SSC-A (B left) and CD11b (B right), proportion of mouse eosinophils that are positive for CD63 (C left) and β2 integrin (C right) from eosinophils alone and upon co-culture with AKP and AKPS tumor organoids (Eos n = 6, Eos^AKP^ and Eos^AKPS^ n = 5 samples). (**D** and **E**) *Spp1* gene expression relative to *Gapdh* (D) and (E) the proportion of double positive (CD206 and CD13) macrophages of murine bone-marrow derived macrophages alone (Mac) and conditioned with AKPS-sup (supernatant) or TGFβ+IL-10 (n = 5 samples each). (**F**) Bubble plot displaying the average gene expression of *Il10* and *Tgfβ1* within annotated mouse scRNAseq data (WT control and tumor n = 5, PHIL control and tumor n = 2, WT peritoneal mets n = 3, PHIL peritoneal mets n = 6 samples). (**G**) *Il10* (left) and *Tgfβ1* (right) gene expression relative to *Hprt* from WT and PHIL whole tumor AKPS tissue (WT n = 10, PHIL n = 7 samples). (**H-J**) CD206^+^ CD13^+^ subsets (H) and representative flow cytometry plots (I), and LAP (TGF-β1)^+^ (J) subsets as a proportion of total macrophages upon preconditioning with TGFβ+IL-10 and additional co-culture with eosinophils (Eos) with and without pretreatment with AKPS supernatant (n = 5 samples each). (A-E, G, H, J) Unpaired t-test for groups of 2, and One-Way ANOVA for groups of 3. Mean ± standard deviation is indicated; stars indicate significance of the p-value (* < 0.05, ** < 0.01, *** < 0.001, **** < 0.0001, ns non-significant).

To explore the molecular mechanisms underlying eosinophil activation, we examined the role of SoxC transcription factors, known regulators of oncoembryonic programs reactivated in CRC (*19*). These transcription factors orchestrate embryonic-like pathways that are co-opted by tumor cells to drive growth, immune modulation, and metastasis (*19*). Notably, co-culture of eosinophils with SoxC-deficient AKPS organoids significantly reduced the proportion of A-Eos compared to WT AKPS organoids (**fig. S5D**). Further analysis of cytokine-regulated gene signatures using Cytokine Signaling scores (CytoSig) (*26*) highlighted IL-33, HMGB1, IL-10 and TGF-β3 as potential mediators of eosinophil activation in both species (**fig. S5E**). The alarmins IL-33 and HMGB1 effectively promoted the maturation of eosinophils into the A-Eos subset *in vitro* (**fig. S5F**). While *Hmgb1* was ubiquitously expressed in the TME, *Il33* expression was restricted to non-immune cells such as fibroblasts, endothelial and epithelial cells (**fig. S5G**). Together, these results underscore the role of tumor-intrinsic factors in modulating eosinophil activation states and functional properties within the TME.

We next investigated factors driving the differentiation of SPP1^+^ macrophages *in vitro*. Although AKPS-sup induced upregulation of the *Spp1* transcript in bone marrow-derived macrophages (**Fig. 5D**), it did not significantly increase the expression of the surface markers CD206 and CD13, which have been reported to reliably identify SPP1^+^ TAMs (*12*) (**Fig. 5E**) and are specifically upregulated in macrophages from AKPS tumors compared to normal adjacent tissue (**fig. S5H**). In addition, the corresponding transcripts (*Mrc1* for CD206 and *Anpep* for CD13) were expressed within the SPP1^+^ TAM cluster in our mouse sequencing dataset (**fig. S3A**). Screening various cytokines revealed that IL-10 and TGF-β strongly induced the SPP1^+^ TAM phenotype both transcriptionally, with upregulation of *Spp1, Fn1*, and *Arg1* (**fig. S5I, J**) and at the protein level (**Fig. 5E**). In contrast, the differentiation of SPP1^+^ TAMs was independent of cytokines such as GM-CSF, IL-4, IL-13, TNF-α and IFN-γ (**fig. S5I)**. Since AKPS organoid supernatant alone did not significantly promote the SPP1^+^ phenotype, the differentiation of this TAM subset may require the presence of additional components beyond tumor-derived factors. Supporting this, while multiple immune and non-immune cell types expressed *Tgfb1*/*TGFB1* in our scRNA-seq datasets, *Il10/IL10* expression was predominantly restricted to the monocyte/ macrophage immune compartment (**Fig. 5F and fig. S5K**). Moreover, both *Tgfb1* and *Il10* transcripts were significantly upregulated in AKPS tumors from PHIL mice compared to their wild-type littermates (**Fig. 5G**), consistent with the higher proportions of SPP1^+^ TAMs observed in these mice.

To explore the relevance of eosinophil-macrophage interactions, we co-cultured TGF-β- and IL-10-conditioned macrophages with eosinophils. Remarkably, the presence of eosinophils was su?cient to strikingly reduce the frequency of CD206^+^ and CD13^+^ macrophages, regardless of their prior preconditioning with AKPS-sup (**Fig. 5H, I**). Furthermore, the expression of LAP (TGF-β1), a marker associated with the pro-tumorigenic activity of macrophages (*27*), was markedly decreased in the presence of eosinophils (**Fig. 5J**). These findings suggest that the differentiation of SPP1^+^ TAMs relies on IL-10 and TGF-β signaling, a process which can be effectively counteracted by eosinophils.

## Discussion

Eosinophils are associated with improved prognosis in several cancer types, particularly those of the GI tract, where their densities have prognostic relevance (*13, 28*). In this study, we present a comprehensive analysis of eosinophil dynamics within the TME, integrating scRNA-seq data from human CRC biopsies and experimental mouse models. Our findings reveal a conserved activation pattern and transcriptional program in tumor-infiltrating eosinophils across species, emphasizing their functional reprogramming within the TME. Notably, we identify and characterize a previously unrecognized eosinophil– macrophage axis that shapes the tumor immune network and plays a pivotal role in CRC progression and metastasis.

Recent advances in scRNA-seq have significantly enhanced our understanding of the complexity of myeloid cells within the TME (*12*, *22*, *23*). However, granulocytic cells like eosinophils often evade detection by classical transcriptomic methods due to their fragile nature and propensity to degranulate, which hampers effective RNA capture. Using the BD Rhapsody platform, which enables higher detection rates of granulocytes (*14*, *29*–*31*), we successfully profiled eosinophils at the single-cell level from CRC patient biopsies, including tumors, normal adjacent tissue and healthy blood samples. Our analysis identified two distinct eosinophil clusters that closely mapped the profiles of our previously described A-Eos and B-Eos murine subsets (*14*).

Our data suggest a progressive activation of blood eosinophils as they transitioned to intestinal tissues and tumors, reflecting their adaptation to the local tissue and tumor microenvironment. Notably, this activation appears to be driven by soluble tumor-derived factors and is influenced by the mutational stage of tumor cells. SoxC transcription factors – key regulators of oncoembryonic programs (*19*)– and their downstream effectors, such as HMGB1, collectively promote the emergence of the activated A-Eos phenotype, underscoring their pivotal role in shaping immune dynamics within the TME.

Activated eosinophils exhibited distinct transcriptional profiles, including genes associated with NF-κB, TNF-α, and IFN-γ signaling, alongside multiple cytokine and chemokine genes. These findings align with a prior study demonstrating the anti-tumor activity of TNF-α/IFN-γ–activated eosinophils in the metastatic lung (*32*). Interestingly, in non-metastatic settings, eosinophils have been shown to control primary tumor growth through direct cytotoxicity or recruitment and activation of T cells (*13, 33*–*36*). In contrast, our findings in the AKPS model reveal a distinct role for eosinophils in constraining peritoneal metastases, without consistently impacting primary tumor size. This discrepancy underscores the likely evolving role of eosinophils during CRC progression, transitioning from localized anti-tumor functions in early-stage disease to limiting dissemination in advanced stages.

A key finding of our study is the role of eosinophils in regulating SPP1^+^ TAMs, a pro-tumorigenic macrophage and subset pro-metastatic implicated in angiogenesis, EMT and immune exclusion (*11, 12, 22*). Interactions between SPP1^+^ TAMs and cancer-associated fibroblasts have been proposed to contribute to the formation of a desmoplastic structure, limiting immune cell infiltration into the tumor core (*37*). This dense, fibrotic matrix may also hinder eosinophil recruitment in advanced CRC, explaining their reduced numbers in later stages of the disease. In both human biopsies and the murine AKPS model, SPP1^+^ TAMs were enriched within tumors but nearly absent in normal adjacent or control tissue, consistent with their *in situ* differentiation from monocyte-like precursors (*22, 23*). In eosinophil-deficient PHIL mice, SPP1^+^ TAMs accumulated in primary tumors, coinciding with increased peritoneal metastases and a mesenchymal phenotype, as well as upregulation of angiogenesis and tissue remodeling genes like *Vegfa* and *Mmp9*. Notably, the proportion of SPP1^+^ TAMs in peritoneal metastases was unchanged in PHIL mice, indicating that eosinophils primarily influence TAM differentiation in primary tumors and act early in the metastatic cascade to prevent invasion and EMT.

Our ligand-receptor interaction analysis reveals significant crosstalk between eosinophils and SPP1^+^ TAMs within TME. Eosinophils may counteract the tumor-supportive phenotype of SPP1^+^ TAMs through interactions involving ICAM-1 and ITGAM/CD11b (*38*). Alternatively, eosinophils may release soluble mediators, such as granular proteins, reactive oxygen species, or pro-inflammatory cytokines like IL-1β or TNFα (*39*–*41*). Additionally, eosinophils may preferentially promote the differentiation of C1QC^+^ macrophages – an alternative subset derived from tumor-infiltrating monocyte-like precursors that could compete with SPP1^+^ TAMs, altering the immune balance within the TME (*23*). Further studies are needed to fully elucidate the mechanisms underlying eosinophil interactions with TAMs, and their role in metastatic progression.

The prolonged progression to metastatic CRC raises the question of whether eosinophils become systemically activated during this process. While blood eosinophils from eosinophilic esophagitis (EoE) patients were transcriptionally similar to those in healthy individuals (*42*), disseminating tumor cells in metastatic CRC may induce systemic transcriptional changes in eosinophils, altering their function in the bloodstream and in distal organs. Larger patient cohorts are required to investigate these dynamics and their implications for CRC progression. However, the fragile nature of eosinophils limits their detection in bulk RNA-seq datasets from public repositories such as TCGA (*43*), emphasizing the need for more refined methodologies.

In summary, our study highlights the critical role of eosinophils in limiting tumor dissemination in a model of advanced CRC. The conserved transcriptional profiles and activation signatures observed in eosinophils across species strongly support the relevance of these findings in CRC patients. While eosinophil-targeting drugs such as mepolizumab and benralizumab effectively reduce eosinophil levels for severe asthma management (*44, 45*), the long-term effect of eosinophil depletion on cancer risk, particularly in tumors with prolonged latency such as CRC, remains uncertain (*46*). Future therapeutic strategies aimed at selectively activating or recruiting eosinophils to modulate the TME may open new avenues for controlling CRC progression and metastasis.

## Materials and Methods

### Mice

C57BL/6J (stock no. 000664) and C57BL/6J-ApcMin/J (stock no.002020) mice were obtained from The Jackson Laboratory. Eosinophil-deficient mice [PHIL (*25*)] were obtained from m J.J. Lee (Mayo Clinic, Phoenix, AZ). C57BL/6NRj mice were purchased from Janvier. C57BL/6J mice were used for AKPS organoid injections and non-tumor bearing control mice, PHIL mice were used as the eosinophil-deficient counterpart. All mice were 8-10 week old males at the start of the experiment. *Il5* transgenic (*Il5tg*) mice were described previously (*47*), and were used to magnetically sort eosinophils for *in vitro* cell culture experiments. Mice were maintained on a 12h light and 12h dark schedule, and water and chow were provided *ad libitum*.

### Animal experimentation

All mouse experiments were performed under the approval of the Cantonal Veterinary Office of Zurich, Switzerland and in accordance with Swiss guidelines.

#### Colonoscopy submucosal injection of AKPS organoids

This procedure was adapted from a prior published paper (*48*). Upon culturing and passaging of AKPS organoids in Matrigel (Corning) *in vitro*, the organoids were mechanically dissociated, and the equivalent of one 40µl Matrigel dome was resuspended in 50µl OptiMEM (Gibco) to equate to one injection in one mouse. 8-10 week old male mice of a given strain were first anesthetized with isoflurane and placed facing up on a 37°C heating pad. Warm PBS was used to flush fecal content from the colon, before the AKPS organoid solution was injected carefully below the mucosa, using a custom needle, syringe and colonoscope. Observation of a visible injection bubble was noted as an indicator for successful injection in the correct location. The mice were then regularly monitored according to the license, until the experimental or humane endpoint.

#### *In vitro* experimentation

##### Organoid culturing

AKP [*Apc*-LOF (loss of function), *p53*-LOF, *Kras*-GOF (gain of function)] (*19*), AKPS [*Apc*-LOF (loss of function), *p53*-LOF, *Kras*-GOF (gain of function), *Smad4*-LOF] (*19*) organoids or AKPS SoxC knockout organoids (*19*) were cultured in Matrigel (Corning) domes of 50 μl, as previously detailed (*49*). The domes were surrounded with complete medium, composed of Advanced DMEM/F12 (Life Technologies) supplemented with 10 mM HEPES (Gibco), 1X GlutaMax (Gibco), 1% penicillin–streptomycin (Gibco), 1X B27 supplement (Gibco), 1X N2 supplement (Gibco) and 1 mM N-acetylcysteine (Sigma-Aldrich). Every 2-3 days, the organoids were split by mechanical dissociation and TrypLE (Gibco, 12604013).

##### Isolation and culture of splenic eosinophils

Spleens of IL-5-tg mice were passed through a 40-µm cell strainer, using a syringe plunger. Ice-cold distilled water was used for red blood cell lysis. Eosinophils were then isolated using anti-SiglecF microbeads (Miltenyi Biotec, 130-118-513), according to manufacturer’s instructions.

##### Isolation and culture of murine BM-derived macrophages

To isolate bone marrow cells, the femur and tibia of mice were flushed using a 23-gauge needle with complete RPMI medium (Gibco, 11875093), and filtered through a 40-µm cell strainer, followed by a red blood cell lysis with ice-cold distilled water for 30 seconds. Cells were then seeded at a density of 0.3×106/ml with complete RPMI medium supplemented with 20ng/ml recombinant M-CSF (BioLegend, 576404), at 37°C. On day 3, more M-CSF supplemented medium was added, and then fully replaced at day 6. On day 7, macrophages were fully differentiated.

##### In vitro conditioning with supernatant of tumor organoids and cytokines

Supernatant of AKPS tumor organoids was prepared by dissociating one organoid dome mechanically. They were cultured in 250µl of complete RPMI medium in a humidified incubator with 5% CO_2_, for 24 h at 37 °C. Eosinophils were magnetically sorted from the spleen of IL-5-tg mice, via positive selection using anti-Siglec-F microbeads (Miltenyi Biotec, 130-118-513), according to the manufacturer’s instructions, or differentiated from bone marrow with mouse SCF (100ng/ml, PeproTech, 250-03) and FLT3-Ligand (100ng/ml, PeproTech, 250-31L) and maintained in complete RPMI supplemented with mouse IL-5 (10 ng ml/1, PeproTech, 215-15). Macrophages were differentiated from the bone marrow of mice and maintained in M-CSF (BioLegend, 576404). Cells were seeded in flat-bottom 48-well plates at a density of 200,000 cells per well (200µl) and conditioned for 16h at 37 °C with cell-free AKPS-supernatant (1:5) or the following mouse cytokines at a concentration of 20ng/µl: TGF-β1 (BioLegend, 763102), IL-10 (PeproTech, 210-10), GM-CSF (PeproTech, 315-03), IL-4 (PeproTech, 214-14), IL-33 (PeproTech, 210-33), TNF (PeproTech, 315-01A), IFNg (PeproTech, 315-05).

##### Quantitative PCR (qRT–PCR)

RNA from either whole tumor tissue or from BM-derived macrophages with conditionings, was isolated using the RNeasy Mini kit (74106 QIAGEN). RNA was isolated according to manufacturer’s instructions, including the optional DNase1 digestion step. cDNA was synthesised with Superscript III reverse transcription (18080-044 QIAGEN). CFX384 Touch Real-Time PCR system (Bio-Rad) was used to measure gene expression, with TaqMan Gene Expression Assays (4331182 Applied Biosystems by Thermo Fisher Scientific) : *Spp1* (Mm00436767_m1), *Tgfb1* (Mm01178820_m1), *Il10* (Mm01288386_m1), *Hprt* (Mm03024075_m1), *Gapdh* (Mm99999915_g1). Relative gene expression was compared to either *Hprt* or *Gapdh* expression, using the 2ΔC(t) calculation.

### Flow cytometry

#### Staining

Cells were surface stained in PBS at 4 °C for 30 minutes with fixable viability dye eFluor 780 (Invitrogen, 65-0865-14), and combinations of the following listed antibodies (1:200, all from BioLegend; unless otherwise stated): CD11b BV510 (M1/70, 101263), CD45 BV650 (30-F11,103151), CD80 BV605 (1:100, 16-10A1, 104729), Ly6G Percp-Cy5.5 (1A8, 127616), MHCII AF700 (M5/114.15.2,107622), PD-L1 PE-Cy7 (1:100, 10F.9G2, 124314), SigF BV421 (E50-2440, 552681 BD Biosciences), β2-integrin AF647 (1:200, 101414), CD206 BV711 (141727), CD13 FITC (111005), LAP (TGF-β1) (141405). Samples were acquired on the LSRII Fortessa or Cytek Aurora 5L (Cytek Biosciences)

#### Data analysis

Analysis of flow cytometry data was performed using FlowJo Software. Relative cell frequencies, cell counts, or Mean Fluorescent Intensities (MFIs) were subsequently plotted in graphical formats using GraphPad Prism.

#### Statistical analysis

All statistical analyses were performed using GraphPad Prism. For statistical comparisons of two groups, two-tailed unpaired t-tests were used. For comparisons of more than two groups, a one-way analysis of variance (ANOVA) was used. Differences were considered statistically significant when P < 0.05.

### TMAs and quantification

#### Human tissue microarrays

The microarrays CO1505 and BC05002b were obtained from TissueArray.com. Deparaffinized sections were subjected to antigen retrieval in 2.4 mM sodium citrate and 1.6 mM citric acid, pH 6, for 25 min in a steamer. Sections were washed with PBST (0.1% Tween in PBS) and blocked for 1 h at room temperature in a blocking buffer (5% BSA, 5% heat inactivated normal goat serum in PBST). Slides were incubated overnight at 4 °C with the following primary antibodies (1:100, in blocking buffer): mouse anti-human MBP-1 (BMK-13, MCA5751 Bio-Rad) and rabbit anti-human PD-L1 (E1L3N, 13684S Cell Signaling). After washing three times with PBST, the following secondary antibodies were added (1:400 in blocking solution) to the slides for 1 h at room temperature: AlexaFluor goat anti-rabbit 594 and AlexaFluor goat anti-mouse 647 (Thermo Fisher Scientific). Slides were mounted in Prolog Gold (P36930 Invitrogen) and imaged on a Nikon Ti2-E inverted microscope, equipped with CrestOptics X-Light v3 confocal disk unit, Lumencor Celesta lasers and Photometrics Kinetix camera.

#### Image analysis for quantification

Only intact cores were counted. ND files were imported in Imaris 9.6.0 and spot objects were created in the green (MBP-1) and red (PD-L1) channels separately (estimated XY diameter = 7 μm, estimated Z diameter = 4 um, quality filter > 6). To quantify the co-expression of PD-L1 and MBP-1, the distance of each spot in the green channel to the nearest spot in the red channel was computed. Green spots (eosinophils) with distance to red spots < 4 μm were considered as active eosinophils (A-Eos) (co-expressing PD-L1 and MBP-1). Green spots with distance to red spots > 4 μm were considered basal eosinophils (B-Eos). The active-to-basal ratio was then computed by dividing the number of active by the number of basal eosinophils in each core. To quantify eosinophils percentage in the core, a spot object in the blue (DAPI) channel was created (estimated XY diameter = 7 μm, estimated Z diameter = 4 um, quality filter > 6) and DAPI cells were counted.

### Human sample collection and ethical statement

#### Collection of human biopsies (normal mucosa and tumor)

Adjacent normal mucosa and tumor tissues were collected from CRC patients with informed written consent, and under approval of the Ethics Committee of Basel, EKBB, no. 2019-02118. Fresh tissues were kept in DMEM (Life Technologies) containing 10% FBS, 1% penicillin–streptomycin (P0781 Sigma) on ice, until processing.

#### Blood collection

Blood was collected from healthy donors with informed written consent and under approval of the cantonal ethical committee of Zurich, Switzerland (BASEC-Nr.: 2019-00837).

### Tissue dissociation for scRNAseq

#### Mouse AKPS tumors

Tumors and metastases were collected at ethical endpoints and then a single cell suspension was prepared using the following protocol. AKPS tumors were cut into small pieces and digested in a shaking incubator at 37°C in RPMI 1640 supplemented with 10% FBS and 100 U/ml penicillin/streptomycin (P0781 Sigma), containing type IV collagenase (C5138 Sigma) and 0.05 mg/ml DNase I (10104159001 Roche). A single cell suspension was then obtained by pushing through a 70-µm cell strainer.

#### Colon from non-tumor bearing mice

Colons from non-tumor bearing mice were collected and cleaned, removing mesenteric fat and intestinal contents. The colons were then cut open lengthwise and washed in PBS, before being cut into 1-2 cm pieces. The colonic tissue was then immersed in falcon tubes containing wash buffer (2% BSA, 100 U ml−1 penicillin–streptomycin (P0781 Sigma) and 5 mM EDTA in HBSS) and placed in a 37°C shaking incubator for 2 x 25 min washes. The wash buffer was replaced in between washes. The tissues were subsequently rinsed with PBS, and then digested 37°C for 50 min in RPMI 1640 supplemented medium, containing 15 mM HEPES (H0887 Sigma), 500 U/ml type IV (C5138 Sigma) and type VIII (C2139 Sigma) collagenase, and 0.05 mg/ml DNase I (10104159001 Roche). Cell suspensions were then obtained by passing the digested mix through a 70-µm cell strainer, followed by centrifugation and layering onto a 40/80% Percoll (17089101 Cytiva) gradient. The interphase was subsequently collected and washed in PBS.

#### Blood from non-tumor bearing mice

Blood from non-tumor bearing mice was collected via terminal cardiac puncture. Red blood cells were subsequently lysed using distilled cold water for 30 s.

#### Human CRC biopsies (tumor and NAT)

Fat tissue and visible blood vessels were removed before tissue processing. Fresh normal mucosa and CRC tissue were washed three times with ice-cold PGA solution (HBSS, 1% penicillin–streptomycin (P0781 Sigma). Tissues were then shaken into 25 mL of DTT solution (PGA solution; 0.1M DTT (Thermo Scientific,707265ML) at 300 rpm for 15 min at RT. After removal of the DTT solution, samples were washed twice with PGA solution and incubated twice for 15 min at 300 rpm at 37 °C in EDTA solution (PGA solution; 1M HEPES (H0887 Sigma); 0.5M EDTA). Tissues were cut into small pieces, washed twice with PGA solution and digested with collagenase A (10103578001 Roche) at 0.16 U/mL and DNase I (10104159001 Roche) at 2 mg/mL in complete RPMI 1640 medium (containing 10% FBS and 1% penicillin–streptomycin (P0781 Sigma)) for 45 min at 37 °C in the GentleMACS dissociator (program: 37C_h_TDK_2). After digestion, cells were filtered through a 100 µm strainer and spun at 200g for 3 min.

#### Human blood processing

Blood was resuspended in 10mL PBS and spun at 400g for 5 min.

### Cell enrichment using magnetic beads

For colonic tissues (AKPS mouse tumors, colon from non-tumor bearing mice, peritoneal metastases, human biopsies) cells were subjected to a CD45^+^ enrichment using anti-CD45 microbeads (mouse: 130-052-301, human: 130-045-801, Miltenyi Biotec) according to the manufacturer’s instructions. Blood samples (blood from healthy individuals, blood from non-tumor bearing mice) were positively enriched for eosinophils using a PE anti-mouse Siglec-F antibody (562068, BD Biosciences; E50-2440) and anti-PE microbeads (130-042-401, Miltenyi Biotec) for mice; and MACSxpress Whole Blood Eosinophil Isolation Kit (130-104-446, Miltenyi Biotec) for human. After the enrichment cells were subjected to morphological examination with Trypan Blue before proceeding for scRNA-seq.

### Single cell RNA sequencing - experimental

#### Cell hashing to pool multiple samples

In order to pool multiple samples for sequencing, single cells of individual samples were labeled with sample tags (633781 BD Human Single-Cell Multiplexing Kit, 633793 BD Mouse Single-Cell Multiplexing Kit) following the manufacturer’s instructions. In brief, 10^6^ cells from each sample were resuspended in a staining buffer (PBS with 1% BSA, 1% EDTA) containing the respective sample tag. After an incubation of 20 min at room temperature, cells were washed twice with a staining buffer, using a centrifugation at 400g for 5 min, before resuspension in 1 ml staining buffer for counting. From each sample, approximately 10-20,000 cells from up to 4 sample tags were pooled, resulting in 60-80,000 cells, for single cell capture.

#### Single cell capture, library preparation and sequencing

Pooled samples were centrifuged a last time at 400g for 5 min, before their resuspension in 650 µl BD Sample Buffer supplemented with a cocktail of Rnase inhibitors (1:1,000 SUPERase 20 U µl^-1^, AM2694 Thermo Fisher Scientific; NxGen Rnase Inhibitor 40 U µl^-1^, 30281-2 Lucigen). Cells were then loaded on a BD Rhapsody cartridge and their RNA was captured with the BD Rhapsody Express Single-Cell Analysis System following the manufacturer’s instructions (BD Biosciences) using version 1 beads for the mouse colonic samples and the enhanced beads for the remaining samples. cDNA and sample tag libraries were prepared with the BD Rhapsody Whole Transcriptome Analysis Amplification Kit (633801 BD Biosciences) following the mRNA Whole Transcriptome Analysis (WTA) and Sample Tag Library Preparation Protocol (BD Biosciences). The quality and quantity of the resulting libraries were then analyzed using a Qubit Fluorometer with Qubit dsDNA HS Kits (Q32851 Thermo Fisher Scientific) and a TapeStation 4200 system with HS D5000 tapes and reagents (5067-5592 Agilent Technologies). Paired-end sequencing was performed on NovaSeq6000 and NovaSeqX systems (Illumina) using NovaSeq 6000 SP100 Reagents (100 cycles) and NovaSeqX Reagents. For v1 beads 20%, and for the enhanced beads 1% PhiX were spiked in for sequencing. Sequencing depth was around 20,000 reads/cell for the WTA library and 500-1000 reads/cell for the sample tag library.

### Single cell RNA sequencing - data analysis

#### Pre-processing, normalization and clustering

Raw sequencing data was demultiplexed using Bcl2fastq (v2.20.0.433, Illumina). Resulting Fastq files were then uploaded to the Seven Bridges Genomics platform and further processed using the BD Rhapsody^TM^ WTA analysis pipeline (v1.12.1) with default settings for the mouse samples. For the human samples the Exact Cell Count parameter was set to 100,000 to increase the chance of identifying eosinophils which often do not get identified because of their low number of expressed genes. For gene mapping a previously generated STAR index with gene code GRCm38 v25 for mouse and GRCh38 v43 for human data was generated using STAR (*50*) (v2.5.2b). Downstream analysis was done in R (v4.3) using the Seurat package (v5.1.0) (*51*) if not specified otherwise. Cells with fewer than 200 gene features, more than 5000 gene features (only for mouse data, no upper cutoff for human data) and more than 25 % of genes mapped to mitochondrial genes were removed. Additionally, cell free RNA contaminations were estimated and removed using the bioconductor package decontX (*52*) (v1.0.0, https://github.com/campbio/decontX), which applies a Bayesian model. All samples were log normalized and scaled using the Seurat functions ‘NormalizeData’ and ‘ScaleData’ while regressing out nFeature_RNA, nCount_RNA and percent.mt. The top 2,000 variable features were used for principal component analysis (PCA) which was followed by the shared nearest neighbor clustering using the Louvain algorithm with multi level refinement. The clustering was then visualized by UMAPs. First clusters were analyzed at low resolutions between 0.1 - 0.5 for broad cell type annotations followed by sub-clustering of individual clusters using the Seurat Function ‘FindSubClusters’ to annotate cellular subtypes at resolutions between 0.1 - 0.5.

#### Integration of datasets

Colonic samples were combined per species and blood samples from healthy individuals were pooled separately. In all cases, the data was integrated using the fast mutual nearest neighbors (FastMNN) approach (‘FastMNNIntegration’ Seurat function). The blood data from non-tumor bearing mice was clustered separately without any batch effect correction. Eosinophils from the human data (colonic and blood derived) were extracted from the annotated data and re-integrated (FastMNN) and re-clustered. Eosinophils from the mouse data were also extracted from annotated data, however, in this case single eosinophils were integrated with a previously published reference (*14*) (non-tumor bearing *Il5-*transgenic mice; eosinophils from colon, blood and bone marrow) using the label transfer approach from the Seurat package. Thereby, the query cells were projected onto the reference UMAP of annotated cells whose label was then transferred to the query cells.

#### Cell type annotation and their comparison between conditions

In all cases clusters were removed if their number of gene features was very small with simultaneously high percentage of genes mapped to mitochondria (low quality), or if they expressed genes from various different cell types (mixed). For automatic annotation the package SingleR (*53*) (v2.4.1, https://github.com/dviraran/SingleR) was used with the murine ImmGen (*54*) (Immunological Genome Project) and the human NovershternHematopoieticData (*55*) data as references, loaded via the celldex (*53*) (v1.12.0, https://github.com/SingleR-inc/celldex) package.

*Mouse CD45*^*+*^ *data:* Clusters were first annotated broadly using the automated cell type annotation approach. Subtypes were then annotated based on known marker genes, their DEGs and the following references: monocytes/macrophages (*56*), T cells (*57*) and ILCs (*58*). Additionally, DEGs were cross-checked in the PangloaDB (*59*) database webtool (https://panglaodb.se).

##### Mouse blood data

Cell clusters were annotated based on automated annotation and investigation of their DEGs.

##### Mouse eosinophilic data

Blood and colonic eosinophils were annotated based on the resulting predicted label from label transfer.

##### Human CD45^+^ data

Clusters were annotated based on DEGs, known marker genes and previously published single cell data references from human cancers (*12, 60, 61*). The annotation of the eosinophils was confirmed using a module score computation approach (see details in a later section ‘Module score computation approach’).

##### Human blood data

Cell clusters were annotated based on DEGs and automatic annotation results. The annotation of the eosinophils was confirmed using a module score computation approach (see details in a later section ‘Module score computation approach’).

##### Human eosinophilic data

Eosinophil subtypes were annotated based on the module score approach (see details in a later section ‘Module score computation approach’).

##### Statistical analysis to assess differences in cell type proportions between conditions

Only CD45 positive cells were used for this analysis. Statistical differences between conditions were estimated using the R package scProportionTest (*62*) (v0.0.0.9000, https://github.com/rpolicastro/scProportionTest).

Thereby, a permutation test was applied to compute p-values, which were further corrected using Benjamini-Hochberg (FDR, false discovery rate). The magnitude of enrichment or depletion for each cell type was calculated via bootstrapping returning a confidence interval (log2FD, log2 fold difference). Changes between conditions were significant if FDR p-values were smaller than 0.05 and the log2FD was larger than 0.5 or smaller than -0.5.

#### Differential gene expression (DEG) analysis

For DEG analysis the Seurat functions ‘FindAllMarkers’ and ‘FindMarkers’ with min.pct = 0.25 and logfc.threshold = 0.25) were used on normalized data. Thereby a non-parametric Wilcoxon Rank-Sum test with Bonferroni correction was applied.

#### Gene set enrichment analysis (GSEA)

DEGs between conditions following the approach explained above were used for gene set enrichment analysis (GSEA) using the R package FGSEA (*63*) (v1.28.0, https://github.com/ctlab/fgsea). To pre-rank, DEGs were sorted in decreasing order using the following formula -log10(p-val_adj)/sign(avg_log2FC), thereby taking the adjusted p-value and the log2 fold change (log2FC) into account. Via null hypothesis testing for each gene ontology (GO) term a normalized enrichment score was calculated and p-values were adjusted using the Benjamaini Hochberg (BH) approach. The genes for the GO terms were extracted from the following Molecular Signature Databases (MSigDB) (*64*, *65*): Reactome, Hallmarks, Kyoto Encyclopedia of Genes and Genomes (KEGG) and Biological Process (BP).

#### Analysis of regulon activity with SCENIC

SCENIC (*66*) (v1.3.1, https://github.com/aertslab/SCENIC) was applied between clusters from the human eosinophilic dataset using default parameters with the following approach: filtering of the gene expression matrix [minCountsPerGene = 3*.01*ncol(exprMat); minSamples = ncol(exprMat)*.01], computation of correlations with the function ‘runCorrelation’, transcription factor target inference using GENIE3 (*67*) (v1.24.0, https://github.com/aertslab/GENIE3) and calculation of gene regulatory networks (GRNs) and their activity scores for each cell with AUCell (*66*) (v1.24.0, https://github.com/aertslab/AUCell).

#### Comparison of mouse and human eosinophils

To quantify the number of shared and unique genes between mouse and human eosinophils all genes with counts larger than zero were counted and compared in a Venn diagram using the R package ggVennDiagram (*68*, *69*) (v1.5.2). Genes where no orthologues were found between the species were dropped and not considered.

#### Module score computation approach

This approach was used to compute the ‘MHC-I and antigen processing’ score, the general eosinophilic and eosinophilic subtype scores. For the general score the top 50 DEGs expressed in eosinophils from annotated colonic mouse cells and for the subtype score the top 50 DEGs between subtypes from wild type eosinophils of the tumor, colon and blood (non-tumor bearing mice) were used. The genes were converted to human gene symbols using the ‘convert_orthologs’ from the orthogene (*70*) Bioconductor package (v1.8.0) and used for the score calculation. For the ‘MHC-I and antigen processing score’ genes from the gene ontology term GO:0002474 were used (mouse: *H2-D1, H2-Q7, H2-K1, H2-T23, H2-Q4, Tap1, Tapbp, B2m, Psmb8, Psme1, Psmb9, Calr, Psmb10, Ncf1, Fcer1g*; human: TAP1, TAPBP, B2M, PSMB8, PSMB1, PSMB9, CALR, PSMB10, NCF1, FCER1G, HLA-A, HLA-C, HLA-B). The module scores were calculated using the Seurat function ‘AddModuleScore’ where the average expression of the module genes was subtracted by the aggregated expression of random background genes. Significant differences between conditions were then identified using a two-sided Wilcoxon Rank-Sum test (function ‘wilcox.test’ from the stats R package).

#### Prediction of ligand-receptor interactions with CellPhoneDB

The Python package CellPhoneDB (*24*) (v5, https://github.com/ventolab/CellphoneDB) was applied to the tumor mouse and human data using Python v3. As inputs the gene count matrix and a metadata file containing the cell type annotations were used. The mouse gene symbols were converted to human symbols because only these can be detected with the CellPhoneDB algorithm. For the conversion the biomaRt (*71, 72*) (v2.58.2) R package was used. For each species we run CellPhoneDB separately using the ‘cpdb_statistical_analysis_method’ with default parameters. In short, for each combination of cell clusters the mean expression of ligands and receptors that are present in the CellPhoneDB repository was calculated. To statistically test their significance a null distribution of means was created from randomly permuted cluster labels. To calculate the total number of interactions from eosinophils and any other cell type we calculated the sum of all interactions going both directionalities (ligand - receptor and receptor - ligand) using the mean values. Mean values were also used to plot L-R interactions between eosinophils and SPP1^+^ TAMs.

#### CytoSig analysis

The average log fold change from significant DEGs (p value adjusted ≤ 0.05, average log2FC > 0.25) between two conditions were used for the web based tool CytoSig (https://cytosig.ccr.cancer.gov/) (*26*). This algorithm was using a two-sided Wilcoxon Rank-Sum test with Benjamini Hochberg p-value adjustment to calculate the predicted response to signaling molecules (chemokines, cytokines and growth factors). The CytoSig database only contains human gene symbols, therefore the mouse gene symbols first needed to be converted using the biomaRt (*71, 72*) (v2.58.2) R package for the mouse data.

#### Plotting

For plotting ggplot2 (*73*) (v3.5.1), pheatmap (v1.0.12, https://github.com/raivokolde/pheatmap) and ComplexHeatmap (*74*) (v2.18.0) were used.

### Schematic drawings

Schematic drawings were generated with Biorender.com.

## End notes

## Acknowledgments

We thank G. De Lange, J. King, M. Nater, V. Marteau, I. Gonzalez-Perez, E. Roussel, P. Krebs, C. Schneider, M. van den Broek and A. Müller for technical support and ideas; E. A. Jacobsen and the Mayo Clinic for providing PHIL mice. We also thank M. Böni for naive blood collection and D. Liberati for helping provide the human samples. We also extend our gratitude to Konrad Basler for his support and for providing the resources necessary for the development of the mouse tumor model. We are also thankful to the Functional Genomic Center Zurich and the Laboratory Animal Services Center (LASC) of the University of Zurich for their help.

## Author contributions

D.R., A.G., T.D., M.B., D.C.,T.V. and H.F. performed the experiments and analysis. A.G., D.R., K.H. and I.C.A. designed the experiments. C.E., C.M., M.S., S.P., T.V. and H.F. provided critical resources and materials. K.H., D.R. and I.C.A. wrote the manuscript. I.C.A conceived and supervised the project.

## Data and materials availability

All data associated with this study are present in the paper or the Supplementary Materials. scRNA-seq data have been submitted to GEO under accession number GSE282765. The analysis code used for this study can be found in the GitHub repository: https://github.com/Arnold-Lab-UZH/Eosinophils_in_CRC.

---

**Supplementary Fig 1.**
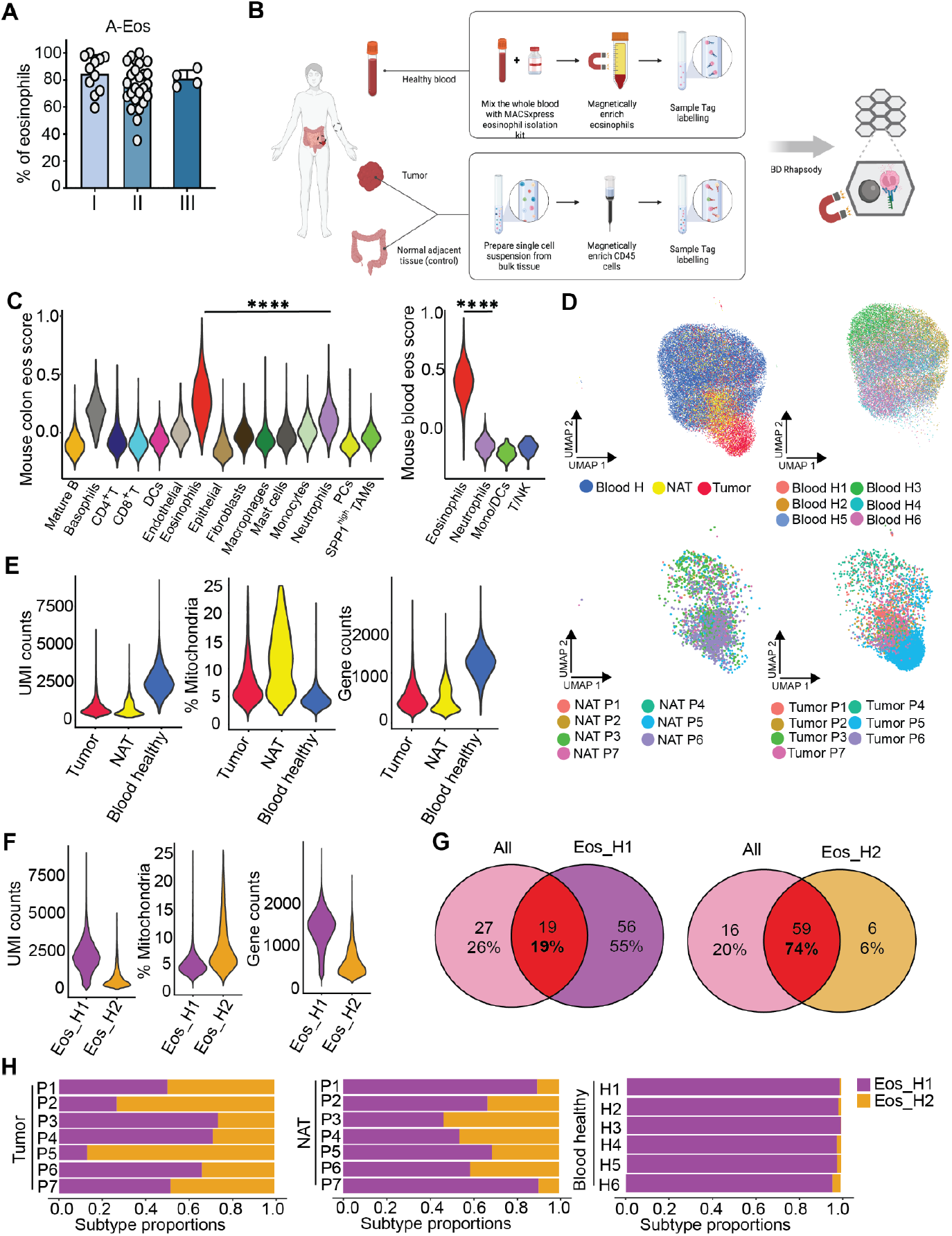
Characterisation of human eosinophils in colorectal cancer at single cell resolution. (**A**) A-Eos frequencies identified on human tissue microarrays via co-localisation of MBP-1^+^ PD-L1^+^ signal, as a proportion of total eosinophils (MBP-1^+^) across CRC stages (I n = 11, II n = 25, III n = 4). (**B**) Schematic drawing of experimental workflow. Created with Biorender.com. (**C**) Violin plots highlighting the mouse eosinophil score on human annotated clusters [left: colon (tumor and NAT data), right: healthy blood data]. Eos: eosinophils, TAMs: tumor associated macrophages, Mono: monocytes, PCs: plasma cells, DCs: dendritic cells, T: T cells, B: B cells, NK: natural killer cells. P-values were calculated using a two-sided Wilcoxon Rank-Sum test between eosinophils and neutrophils. (**D**) UMAP of integrated human eosinophils displaying different tissues/samples. Blood H: blood healthy. (**E** and **F**) Quality control measurements [unique molecular identifier (UMI), percentage of mitochondria to cytoplasmic genes (% mitochondria) and gene counts] between tissues (E) and eosinophil clusters (F). (**G**) Venn diagrams comparing the number of significant DEGs (differentially expressed genes) upregulated in the tumor compared to NAT eosinophils between all eosinophils to only Eos_H1 (left) or Eos_H2 (right) respectively. Non-parametric Wilcoxon Rank-Sum test and Bonferroni correction of p-values. Genes considered significant if p-value adjusted ≤ 0.05 and Log2 Fold Change > 0.25. (**H**) Barplots showing eosinophilic subtype proportions between different patients (left: tumor, middle: NAT) and Blood healthy (right). (C-H) Tumor and NAT n = 7, blood healthy n = 6 samples. NAT: normal adjacent tissue.

**Supplementary Fig 2.**
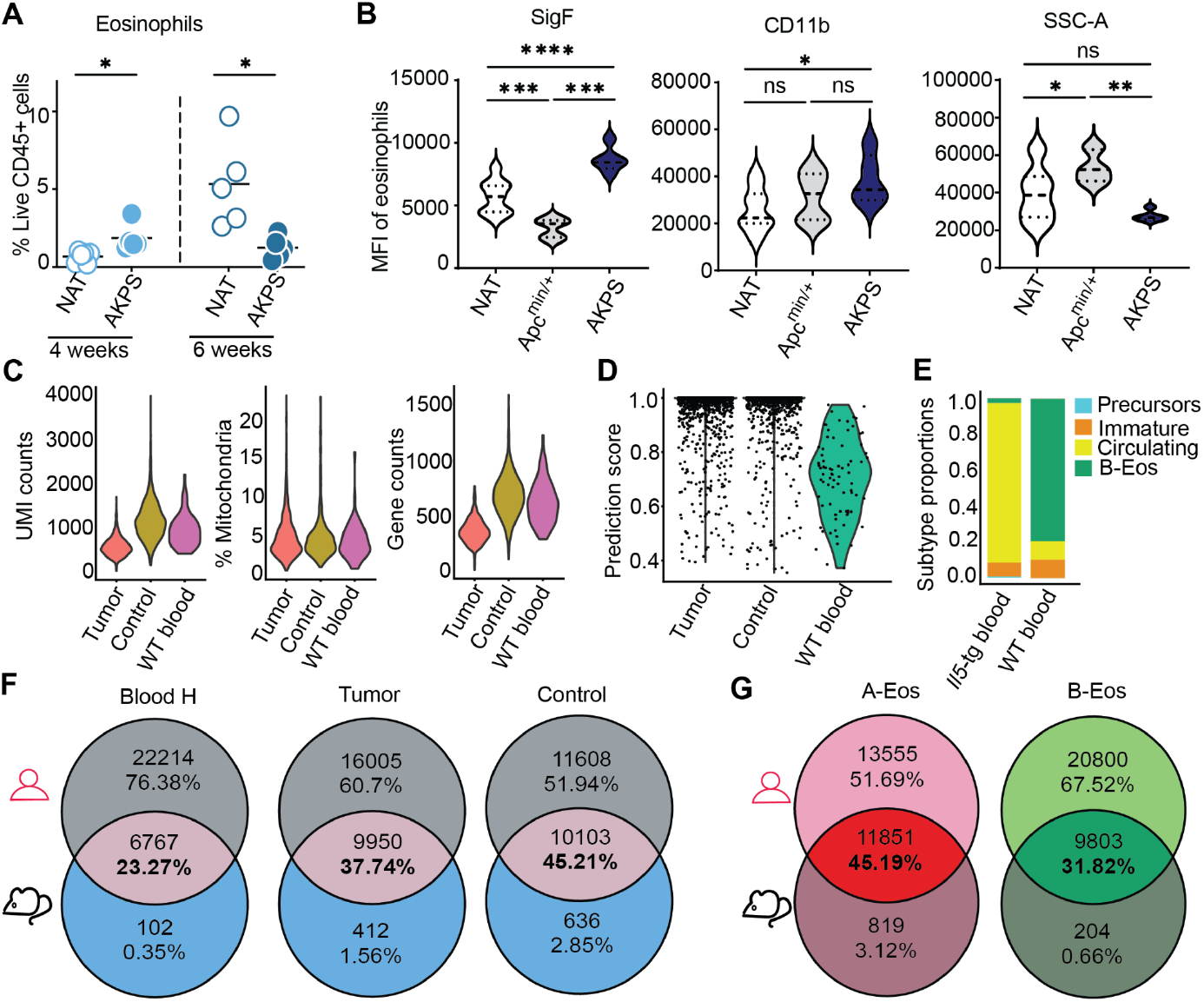
Analysis of mouse CRC eosinophils and their comparison to the human counterparts. (**A**) Frequency of eosinophils from CD45^+^ immune cells in NAT and AKPS tumors after 4 and 6 weeks (NAT and AKPS 4 weeks n = 6, NAT and AKPS 6 weeks n = 5 samples). Unpaired t-test. (**B**) SiglecF (SigF) (left), CD11b (middle) and SSC-A (right) MFI (Mean Fluorescence Intensity) of eosinophils from NAT, Apc^min/+^ adenoma and AKPS tumor (NAT n = 19, Apc^min/+^ n = 7, AKPS n = 5). One-Way ANOVA. (**C**) Quality measurements of eosinophils from tumor, control and blood from non-tumor bearing mice. Left to right: unique molecular identifier (UMI), percentage of mitochondria to cytoplasmic genes (% mitochondria) and gene counts. (**D**) Violin plot indicating the prediction score calculated by the label transfer algorithm to integrate eosinophils from tumor, control and blood WT (wild type) with the *Il5-*tg (transgenic) reference dataset (*14*). (**E**) Barplot comparing eosinophil subtype proportions between *Il5*-tg and blood WT. (**F** and **G**) Venn diagram revealing the number of shared genes (gene counts > 0) expressed by eosinophils between human and mouse tissue (F) and eosinophils subtypes (G). (A, B) Mean ± standard deviation is indicated; stars indicate significance (* < 0.05, ** < 0.01, *** < 0.001, **** < 0.0001, ns non-significant). (C-G) Mouse data n = 5 samples each tissue. (F, G) Human data tumor and NAT n = 7, blood healthy n = 6 samples. (A, B, F, G) NAT: normal adjacent tissue.

**Supplementary Fig 3.**
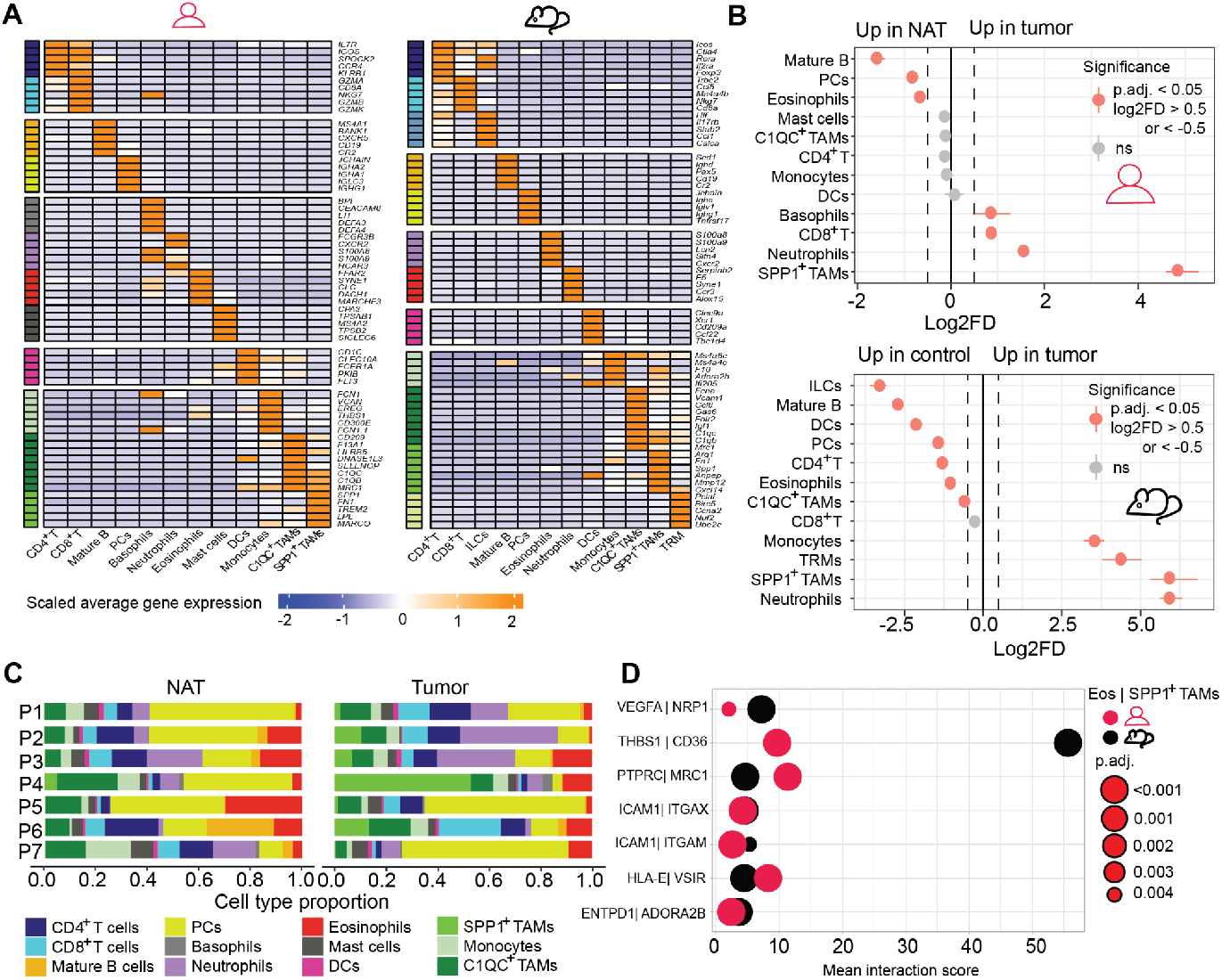
Analysis of immune cells of the tumor microenvironment in human and mouse colorectal cancer. (**A**) Heatmap indicating the scaled average gene expression of selected marker genes within each annotated cell type cluster (human data: left, mouse data: right). (**B**) Point-range plots of statistical analysis of CD45^+^ cell type proportions between control and tumor (human data: upper, mouse data: bottom). Permutation test; Benjamini-Hochberg adjusted p-values (p.adj); log2FD: log2 fold difference. (**C**) Barpot comparing the CD45^+^ cell type proportions between NAT and tumor from human data. (**D**) Bubble plot depicting the shared significant ligand-receptor interactions between eosinophils (Eos) and SPP1^+^ TAMs. The mean interaction score is shown on the x-axis and the dot size defines the significance (null hypothesis testing). (A-C) T: T cells, DCs: Dendritic cells, TAMs: Tumor associated macrophages, B: B cells, Mast: Mast cells, PC: Plasma cells, TRM: Tissue resident macrophages. (A, B) Mouse data control and tumor n = 5 samples each. (A-D) NAT: normal adjacent tissue. Human data tumor and NAT n = 7 samples each.

**Supplementary Fig 4.**
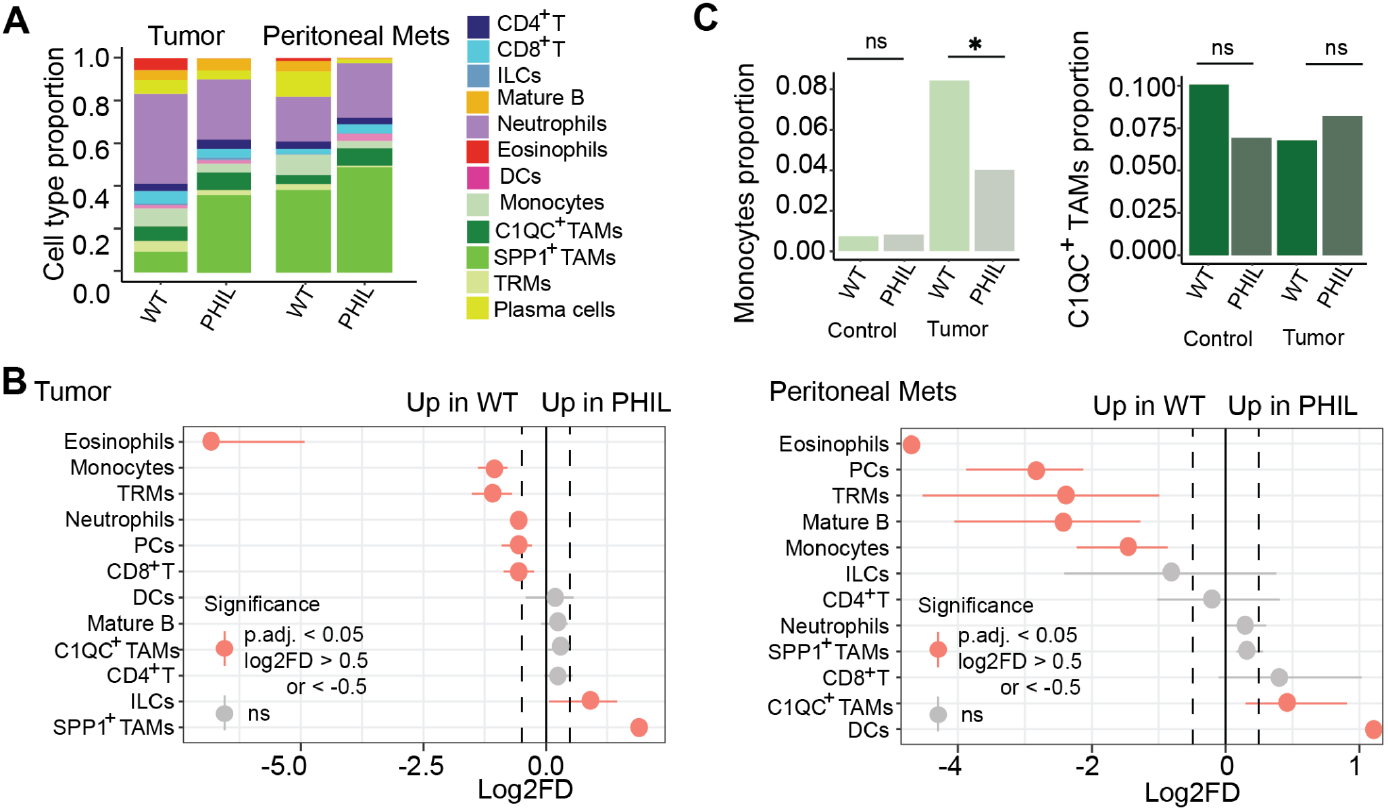
Influence of eosinophil depletion on the tumor microenvironment focusing on SPP1^+^ tumor associated macrophages. (**A** and **B**) Barplots (A) and Point-range plots (B) comparing the cell type proportions between WT and PHIL tumor (left) and peritoneal metastasis (right). P.adj: adjusted p-value. Mets: Metastases. T: T cells, ILCs: Innate lymphoid cells, DCs: Dendritic cells, Mono: Monocytes, TAMs: Tumor associated macrophages, TRMs: tissue resident macrophages, B: B cells. (**C**) Bar plots of Monocytes (left) and C1QC^+^ TAMs (right) comparing the proportions between WT and PHIL from different tissues. Stars indicate statistical significance (* < 0.05, ns non-significant). (C, B) Permutation test, Benjamini-Hochberg adjusted p-values. (A-C) WT control and tumor n = 5, PHIl control and tumor n = 2, WT peritoneal metastases n = 3, PHIL peritoneal metastases n = 6 samples.

**Supplementary Fig 5.**
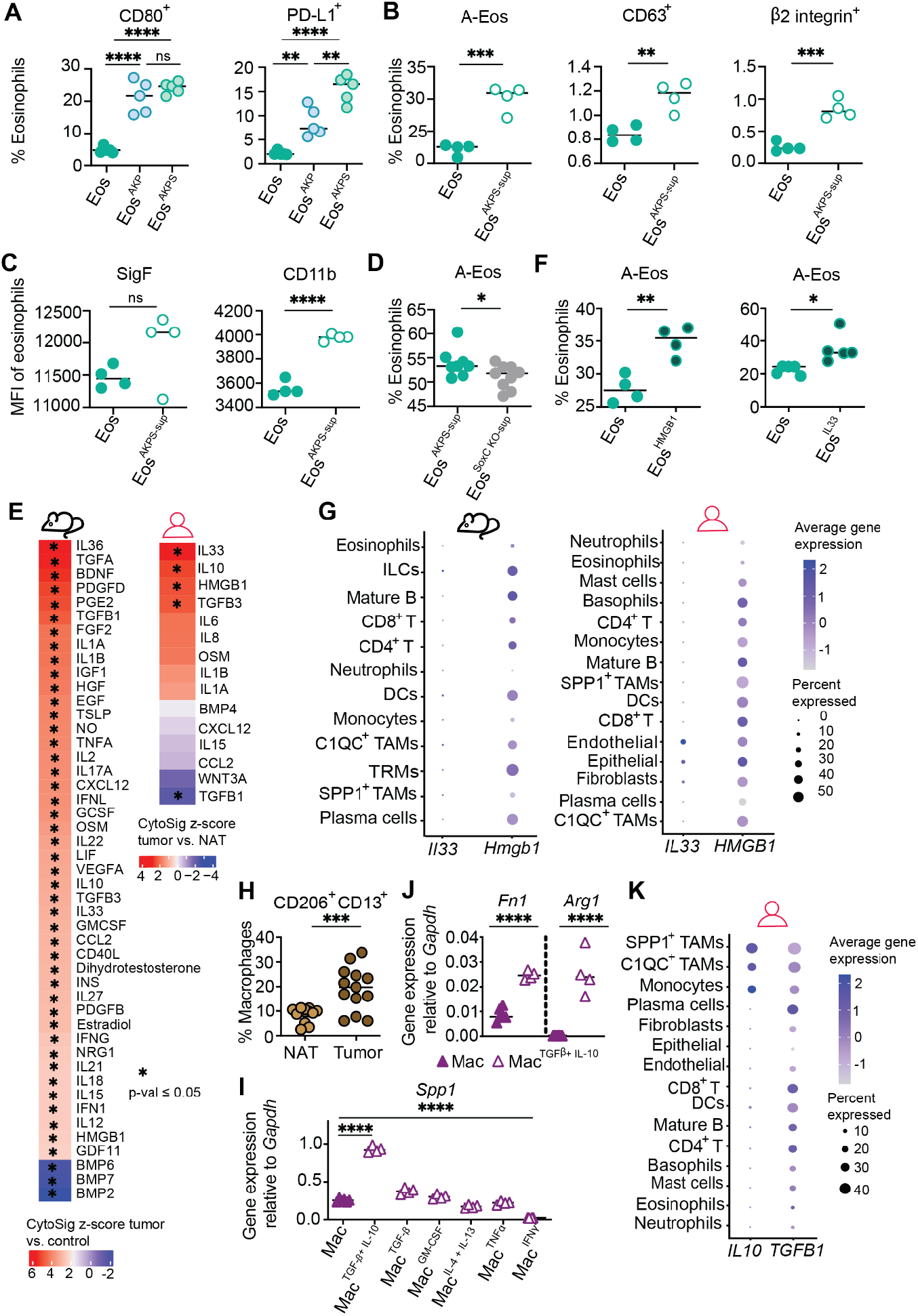
Eosinophils restrict SPP1^+^ macrophage polarization through disruption of IL-10 and TGF-β signaling. (**A**) Proportion of CD80^+^ (left) and PD-L1^+^ (right) splenic eosinophils, upon co-culture with AKP and AKPS organoids (n = 5 samples each). (**B** and **C**) A-Eos subset (B left), CD63^+^ (B middle), β2 integrin^+^ (B right) cells as a percentage of eosinophils, and MFI (Mean Fluorescence Intensity) of SigF (Siglec F) (C left) and CD11b (C right) from *in vitro* culture of eosinophils alone (Eos) and upon conditioning with AKPS-sup (supernatant) (n = 4 samples each). (**D**) A-Eos subset as a percentage of eosinophils upon conditioning with supernatant either from AKPS organoids (AKPS-sup) or SoxC KO AKPS organoids (SoxC KO-sup) (Eos^AKPS-sup^ n = 8, Eos^SoxC KO-sup^ n = 9 samples). (**E**) Heatmaps showing the CyoSig z-score between tumor and control eosinophils within mouse (left) and human (right) datasets. Stars indicate statistical significance (adjusted p-value ≤ 0.05, two-sided Wilcoxon Rank-Sum test, Benjamin Hochberg adjustment). Mouse data control and tumor WT n = 5 samples each. (**F**) A-Eos subset, as a percentage of eosinophils upon conditioning with cytokines HMGB1 (left) or IL-33 (right) (n = 4 samples each). (**G**) Bubble plot showing the gene expression for *Il33/IL33* and *Hmgb1/HMBG1* in mouse (left) and human (right) scRNA-seq data. Mouse WT control and tumor n = 5, mouse PHIL control and tumor n = 2, mouse WT peritoneal metastasis n = 4, mouse PHIL peritoneal metastasis n = 6. (**H**) CD206^+^ CD13^+^ subset of macrophages in NAT (normal adjacent tissue) and tumor (n = 13 samples each). (**I** and **J**) Expression of *Spp1* (I) and *Fn1* and *Arg1* (J) relative to *Gapdh*, in bone-marrow derived macrophages alone (Mac) and conditioned with combinations of cytokine treatments (Mac n = 6, all remaining conditions n = 4). (**K**) Bubble plot showing the gene expression for *IL10* and *TGFB1* in human scRNA-seq data. (A-D, F, H-J) Unpaired t-test for groups of 2, and One-Way ANOVA for groups of 3. Mean ± standard deviation is indicated, stars indicate significance of the p-value (* < 0.05, ** < 0.01, *** < 0.001, **** < 0.0001). (E, G, K) Human data NAT (normal adjacent tissue) and tumor n = 7.

