## Supplemental Tables for "Eosinophils restrict CRC metastasis by inhibiting pro-tumorigenic SPP1^+^ macrophage differentiation"

**Sup. Table 1. Patient information.**

| Sample | Primary Tissue | Tissue Diagnosis (as reported in the surgical pathology report) | Sex | Age at Surgery (Years) | TNM classification: tumor (T), node (N), metastasis (M) | Tumor Grading (WHO) | Treatment received prior to Surgery (1=treated; 0 = untreated) | Type of Treatment | Genetic alteration (MSI status) |
|---|---|---|---|---|---|---|---|---|---|
| P1 | Rectum | Intestinal adenocarcinoma of the rectum | M | 73 | pT3 pN0(0/47) L0 V0 Pn0 R0 | G2 | 0 | | MSS |
| P2 | Rectum | Intestinal adenocarcinoma with infiltration of the muscularis propria | F | 57 | pT2 pN0(0/44) L0 V0 Pn0 R0 | G2 | 0 | | MSS |
| P3 | Rectum | Intestinal adenocarcinoma with infiltration of the muscularis propria | M | 86 | pT2 pN0(0/18) L0 V0 Pn0 R0 | G2 | 0 | | MSS |
| P4 | Rectum | Intestinal adenocarcinoma of the rectum with invasion of ventrally involved soft tissue | M | 58 | ypT4a pN1b(2/24) L1 V1 Pn0 R0 | G2 | 1 | Chemotherapy | MSS |
| P5 | Colon | Intestinal adenocarcinoma of the sigmoid colon with incipient infiltration into the pericolic fat tissue | F | 57 | pT3 pN0(0/35) L0 V0 Pn0 R0 | G2 | 0 | | MSS |
| P6 | Rectum | Intestinal adenocarcinoma with infiltration of the muscularis propria | F | 62 | ypT2 pn0(0/16) L0 V1 Pn0 R0 | G2 | 1 | Neoadjuvant | MSS |
| P7 | Colon | Invasive intestinal adenocarcinoma of the cecum with small focal infiltration into the pericolic fat tissue | | 84 | pT3 pN0(0/21) L0 V0 Pn0 R0 | G2 | 0 | | MSS |

**Sup. Table 2. Significant DEGs tumor versus control (NAT) eosinophils.**

| Increased in tumor eosinophils | | | | Increased in control (NAT) eosinophils | | |
|---|---|---|---|---|---|---|
| **Gene** | **Avg_log2FC** | **P_val_adj** | | **Gene** | **Avg_log2FC** | **P_val_adj** |
| **CCL3L1** | 2.959706829 | 3.37E-81 | | **MT-CO1** | 2.119460447 | 2.23E-308 |
| **CCL3** | 2.886701774 | 1.42E-82 | | **MT-ND1** | 1.976547363 | 9.69E-188 |
| **CCL4L2** | 1.62897846 | 2.14E-58 | | **MT-ND4L** | 1.904551241 | 2.08E-26 |
| **CCL4** | 1.531448417 | 1.01E-49 | | **MT-CYB** | 1.869311825 | 6.64E-221 |
| **CD83** | 1.447589515 | 8.32E-27 | | **MT-CO2** | 1.550382665 | 5.45E-297 |
| **CD69** | 1.379795141 | 6.25E-38 | | **MT-ND4** | 1.538376632 | 6.29E-271 |
| **IGKC** | 1.372125519 | 0.00071804 | | **MT-ND3** | 1.53740318 | 3.68E-232 |
| **PIM2** | 1.302914117 | 2.53E-18 | | **MT-ATP8** | 1.517656723 | 1.86E-18 |
| **ADGRE5** | 1.264038783 | 3.13E-62 | | **MT-ATP6** | 1.373764548 | 2.04E-224 |
| **CEBPB** | 1.141025539 | 6.89E-20 | | **MT-CO3** | 1.348225552 | 8.97E-294 |
| **EGR1** | 1.115279015 | 5.9E-22 | | **MT-ND5** | 1.318461795 | 2.48E-58 |
| **BTG2** | 1.110949213 | 2.2E-50 | | **MT-ND2** | 1.023659854 | 4.92E-168 |
| **PLEK** | 1.096011719 | 3.88E-53 | | **CD55** | 1.009709723 | 5.74E-48 |
| **HLA-DRA** | 1.083151341 | 5.33E-34 | | **RPS24** | 0.837871327 | 1.51E-06 |
| **HSP90B1** | 1.02211432 | 4.78E-14 | | **GLCCI1** | 0.836871315 | 2.85E-21 |
| **CXCR4** | 0.994409499 | 4.22E-44 | | **TMCC3** | 0.72730784 | 1.31E-20 |

| | | | | | | |
|---|---|---|---|---|---|---|
| **CYBB** | 0.96574209 | 2.39E-13 | | **PAK1** | 0.636469043 | 5.62E-10 |
| **TSC22D3** | 0.879054516 | 0.0002197 | | **SCLT1** | 0.61846256 | 5.08E-20 |
| **GK** | 0.86954227 | 1.81E-16 | | **RANBP2** | 0.588778309 | 7.24E-07 |
| **RHOH** | 0.867720607 | 9.86E-12 | | **MAILR** | 0.567389986 | 5.05E-09 |
| **FFAR2** | 0.860696711 | 1.81E-41 | | **PDE3B** | 0.56048707 | 1.35E-20 |
| **TYROBP** | 0.840810441 | 0.000910254 | | **MAP3K8** | 0.557507294 | 1.08E-05 |
| **GPCPD1** | 0.826158687 | 1.64E-15 | | **EEF1A1** | 0.546241745 | 0.002972007 |
| **BIRC3** | 0.819911936 | 3.36E-32 | | **CHD1** | 0.545155822 | 2.12E-21 |
| **RAB11FIP1** | 0.81988108 | 3.57E-24 | | **MT-RNR1** | 0.530674577 | 1.84E-46 |
| **CXCL8** | 0.80298517 | 2.89E-05 | | **MCTP2** | 0.495845321 | 6.73E-11 |
| **GNG2** | 0.776629235 | 4.83E-13 | | **PABPC1** | 0.492638253 | 0.000184663 |
| **SAT1** | 0.752325796 | 9.19E-44 | | **THBS1** | 0.447351667 | 3.43E-39 |
| **G0S2** | 0.748822358 | 1.92E-07 | | **KDM6B** | 0.427491287 | 6.16E-13 |
| **CD44** | 0.710470426 | 3.67E-11 | | **SYNE2** | 0.419678179 | 3.29E-15 |
| **ANXA1** | 0.704160908 | 5.84E-08 | | **PTPRE** | 0.416898056 | 0.000128715 |
| **HLA-C** | 0.667382119 | 3.24E-34 | | **SVIL** | 0.40305672 | 0.018310257 |
| **CAMK1D** | 0.665805234 | 4.17E-15 | | **GPR65** | 0.401524047 | 0.034723251 |
| **CFLAR** | 0.664094669 | 6.7E-09 | | **UTRN** | 0.391259993 | 0.005026032 |
| **FOS** | 0.663445001 | 2.9E-25 | | **SYAP1** | 0.387299962 | 0.00350207 |
| **LYN** | 0.623480805 | 2.61E-26 | | **PTPN12** | 0.386135949 | 0.000484053 |
| **TNFAIP3** | 0.617334181 | 1.53E-27 | | **FGD4** | 0.380140266 | 4.9E-10 |
| **ADGRE2** | 0.59985374 | 1.45E-11 | | **GABPB1** | 0.365871716 | 0.038334395 |
| **DENND4A** | 0.575509487 | 1.09E-19 | | **PSEN1** | 0.36329225 | 0.000703016 |
| **PLIN2** | 0.569345016 | 2.11E-09 | | **EIF1** | 0.354919835 | 8.85E-06 |
| **PELI1** | 0.564676221 | 2.4E-08 | | **BCL2A1** | 0.354592963 | 0.000173515 |
| **NFKBIA** | 0.561237228 | 9.09E-12 | | **PLXNC1** | 0.335155572 | 0.032839625 |
| **RIPOR2** | 0.550962518 | 6.64E-10 | | **PDE4D** | 0.287604758 | 0.000329734 |
| **NR4A3** | 0.550251863 | 0.000574056 | | | | |
| **ZFP36** | 0.540972788 | 2.15E-15 | | | | |
| **NFAT5** | 0.540003462 | 4.99E-09 | | | | |
| **ANKRD28** | 0.539075185 | 1.5E-08 | | | | |
| **TXNIP** | 0.509711349 | 9.88E-08 | | | | |
| **B2M** | 0.495386403 | 2E-30 | | | | |
| **DUSP1** | 0.483291754 | 0.007984933 | | | | |
| **WSB1** | 0.478522243 | 5.46E-07 | | | | |
| **FNDC3B** | 0.464900609 | 2.42E-19 | | | | |
| **RAB1A** | 0.459224542 | 4.24E-08 | | | | |
| **ADGRE1** | 0.457445279 | 1.22E-07 | | | | |

| | | |
|---|---|---|
| **HSPD1** | 0.443327573 | 1.31E-11 |
| **HIF1A** | 0.438696267 | 1.75E-15 |
| **SQSTM1** | 0.430092106 | 3.77E-08 |
| **MCTP1** | 0.416823044 | 0.00059321 |
| **ALOX5AP** | 0.406295427 | 3.18E-12 |
| **ARRB2** | 0.405730804 | 0.00751059 |
| **LINC-PINT** | 0.396371123 | 1.06E-06 |
| **PSTPIP2** | 0.395676433 | 0.000380432 |
| **CDC42SE2** | 0.373328952 | 0.002137169 |
| **VASP** | 0.368963552 | 2.08E-05 |
| **SOD2** | 0.357991221 | 0.002465815 |
| **PICALM** | 0.35471985 | 0.01814821 |
| **RIPK2** | 0.352042022 | 0.000103993 |
| **FTH1** | 0.348035861 | 2.23E-13 |
| **PNISR** | 0.330987785 | 0.040697382 |
| **MCL1** | 0.309348248 | 2.64E-09 |
| **NCOR1** | 0.304899213 | 0.002159727 |
| **SIK3** | 0.275573124 | 0.000539833 |
| **LCP2** | 0.271240264 | 0.001039631 |
| **JUND** | 0.264856718 | 6.41E-07 |
| **TMSB4X** | 0.261407981 | 9.89E-21 |
